# Cell layer-specific modification of cell wall is associated with exo-mesocarp split in pistachio (*Pistacia vera L*.)

**DOI:** 10.1101/2025.06.18.660460

**Authors:** Shuxiao Zhang, Minmin Wang, Shaina Eagle, Alisa Chernikova, Kaleigh Marie Bedell, Phuong Tran, Chaehee Lee, Jaclyn A. Adaskaveg, Yiduo Wei, Rolando Lopez, Annamarie Basco, Phoebe Gordon, Barbara Blanco-Ulate, Grey Monroe, Georgia Drakakaki

## Abstract

Pistachio (*Pistacia vera*) is a drought and salinity-tolerant perennial whose fruit features a fleshy exo-mesocarp, or “hull,” that protects the kernel. Hull development and degradation are key to kernel quality, yet the anatomy and mechanisms driving hull breakdown during late-stage development remain largely unknown.

Here, we show that the hull contains anatomically distinct layers of hypodermal parenchyma and filler parenchyma. Using a combination of transcriptome analyses and immunohistochemistry, we show that changes in pectin associated gene expression and modification of this polysaccharide are involved in hull cell size increase, loss of cell-cell adhesion, and softening.

Anatomical analysis shows that filler parenchyma expands during late-stage hull development while hypodermal parenchyma remains constant in size. Field data suggest that irrigation and humidity affect pistachio hull split, implicating a role for water status in cell expansion. In summary, the complex interplay between molecular, cellular, and environmental changes suggests that cell layer–specific modifications of the cell wall are linked to exo-mesocarp splitting, forming a model for understanding the mechanism of fruit split during ripening in non-berry fruit crops.

**Highlight:** Cell-layer specific modifications of the cell wall are associated with cell expansion and loss of cell-cell adhesion, leading to hull split during late-stage pistachio fruit development.

## Introduction

Fruit cracking, or splitting of the pericarp during ripening, has been modeled in many species including tomato (*Solanum lycopersicum*), strawberry (*Fragaria x ananassa*), and grapes (*Vitis spp.*)(B. M. Chang et al., 2019; B.-M. Chang & Keller, 2021; Correia et al., 2018; Domínguez et al., 2012; Hurtado & Knoche, 2023; La Spada et al., 2024; Michailidis et al., 2020). Fruit cracking during fruit growth and maturation can lead to additional issues such as insect damage, fungal infections, and cosmetic defects, which in turn reduce the marketable yield of the crop (La Spada et al., 2024; Santos et al., 2023). Fruit cracking has been linked to changes in the fruit cell wall that occur during fruit ripening, particularly to modifications in pectin, as well as changes in the fruit cuticle structure. For example, suppression of polygalacturonase (PG), an enzyme involved in pectin degradation, and expansin, proteins known for their role in cell wall loosening, in tomato can reduce epidermal cracking, resulting in firmer fruits with thicker cell walls and cuticle deposition (Cantu et al., 2008; Jiang, Lopez, Jeon, De Freitas, et al., 2019). However, pericarp splitting of other non-berry fleshy fruits is less well studied.

Modification of the cell wall is critical for cell swelling, which is a source of physical tension driving fruit split (Cosgrove, 2000, 2024; Lahaye et al., 2021; Schumann & Knoche, 2020). Cell-cell separation and cell wall swelling can cause the exocarp cells to “unzip”, leading to cracking in the sweet cherry skin (*Prunus avium*), as described in the “zipper model” (Brüggenwirth et al., 2014; Grimm et al., 2019; Schumann & Knoche, 2020). In addition to changes in the exocarp, swelling of the mesocarp from exposure to water when microcracks form on the epidermis is also implicated in fruit cracking (Grimm et al., 2019). In apples (*Malus x domestica*) and sweet cherry, changes in the composition of pectin and hemicellulose in cell walls are implicated in the fruit cell wall’s ability to take up and retain water, which ultimately affects the mesocarp’s texture due to turgor (Lahaye et al., 2021). However, the relationship between cell swelling, cell wall modification, and water status has not been consistently investigated in the context of fruit splitting in many fleshy fruit species.

The difference in stiffness between fruit skin and the fruit interior has also been implicated in fruit cracking. In grapes, epidermal and cuticle stiffness, water status, texture of the fruit and programmed cell death all contribute to maintenance of fruit integrity (B. M. Chang et al., 2019; B.-M. Chang & Keller, 2021; Tilbrook & Tyerman, 2008). The current paradigm for fruits with thin exocarp and thick fleshy mesocarp is that the softening of the fruit, which results in a more liquid-like interior, changes the physical strain placed on the skin, where its mechanical properties, such as its elastic modulus, help determine the amount of strain the skin can withstand before splitting (Bargel et al., 2004; B. M. Chang et al., 2019; Considine & Brown, 1981; Knoche et al., 2004). In contrast, in non-berry fruits with relatively thicker exocarp, thinner mesocarp, heavily lignified endocarps, and large seeds, such as the pistachio, walnuts, and almonds, less is known regarding the process of fruit cracking.

Pistachio, or the fruit of *Pistacia vera,* is a popular woody nut crop and emerging model species due to its nutritional benefits and its relatively high salt and drought tolerance. (Bulló et al., 2015; Ferguson et al., 2002; Godfrey et al., 2019). Nonetheless, irregular weather and the resulting abnormal temperature patterns and water availability can affect the integrity of the pistachio exo-mesocarp, known as the hull, which is critical for protecting the edible kernel from pests and pathogens. During late-stage of pistachio maturation, the hull naturally detaches from the shell and the shell splits along a dorsal-ventral suture while the hull remains intact (Pana hi &Khezri, 2011; Polito & Pinney, 1999). Although the exo-mesocarp (hull) is a critical tissue for fruit integrity and consumer safety (Doster et al., 2001; Doster & Michailides, 1995, 1999), relatively little is known regarding the anatomy of its development under different environmental conditions.

Several distinct hull split phenotypes can occur that impact the economic value and safety of the pistachio. One phenotype is “early splitting”, in which both the hull and the shell “crack” along the suture prior to harvest, leaving the kernel directly exposed to the environment (Doster et al., 2001; Sommer et al., 1986). The hull can also “crack” adjacent to the suture, with the shell still providing some protection to the kernel, although the loss of hull integrity still increases susceptibility to pests and pathogens (Doster et al., 2001). Another category of loss of hull integrity is “tattering”, where the outer layers of the hull become torn and tattered in appearance (Sommer et al., 1986) while the shell and kernel may or may not be exposed. Although the phenotype of hull split has been described in field conditions and is well-known to growers, the cell types involved and mechanisms behind the different phenotypes of hull split are currently unknown. Hull softening occurs during late-stage pistachio fruit development (Adaskaveg et al., 2025) and water status has been implicated in pistachio hull cracking (Doster et al., 2001). It is unknown if these factors can induce cell swelling or if changes in hull texture are key mechanisms driving hull split.

In our study, we used transcriptome-level analysis, complemented with hull anatomy assessment and field data, to identify mechanisms that affect hull integrity in pistachio. Our study identified cell types involved in hull split and their corresponding cell wall modifications, along with candidate genes associated with hull ripening. Furthermore, we showed that the water status of the fruit and cell-type specific cell expansion interact with cell wall modifications, which contribute to the different types of hull split.

## Material & methods

### Plant material and sampling

In 2021, the fruits from trees of *P. vera* ’Kerman’ or ’Golden Hills’ cultivar were sampled from a commercial orchard in Cantua Creek, California. ‘Kerman’ and ‘Golden Hills’ scions are grafted on ’Pioneer Gold I’ (*P. integerrima*) rootstocks, with Kerman trees ∼ 20 years in age and ‘Golden Hills’ ∼ 15 years in age. During the flowering period, 6-12 trees per cultivar were tagged. On each tree, 4 to 6 branches that were spaced out throughout the canopy, each containing only one inflorescence per branch to minimize the effect of fruit load and sun exposure on development and phenology, were selected and tagged for analysis. The end of the flowering period, where between 75 to 80% of the trees were post flowering, was denoted as 0 days post anthesis (dpa). During each time point sampled, one tagged cluster per tree from six trees were harvested and stored at 4 °C in a cooler and transported back to the laboratory for processing, which occurred within 24 h of harvest.

Similarly, in 2022, fruits of ’Kerman’, ’Golden Hills’ and ’Lost Hills’ were tagged in a commercial orchard in Mendota, California. The scions were between seven to eight years in age and were grafted on ‘UCB1’ (*P. atlantica* × *P. integerrima*) rootstock. In 2023, fruits of ’Golden Hills’ were tagged in a commercial orchard located in Woodland, CA, USA. The trees are ∼ 8-9 years in age and grafted on ’UCB1’ rootstock.

In 2024 fruits of ‘Golden Hills’ were sampled from commercial orchards located in two sites in Fresno, California for comparisons between irrigation treatments. Both were mature orchards grafted on UCB-1 rootstock, using double-line drip irrigation. Trial A was managed organically, Trial B was managed conventionally. At each location, the grower’s standard irrigation practices were used as the control (100% applied water), thus the amounts of applied water differ at each location. Irrigation treatments were implemented by altering the number of irrigation lines supplying water. For the 50% treatment, a valve was installed on one of the lines, and was shut off for most of the season. For the 150% treatment, an additional line was installed. To ensure that fertilizer applied through the irrigation system was not altered, the researchers turned off the additional line in the 150% treatment, and turned on the line in the 50% treatment during fertigation events (i.e., during fertilization events, each treatment had two lines supplying water and fertilizer). Each block was three rows wide, and only the central trees in the center row were monitored. This ensured that the trees that were being monitored were not picking up water from a neighboring row. As the treatments were implemented at the irrigation system riser, the number of trees in each row differed: at Trial A, there were 38 trees in each plot row (114 in each experimental plot). At Trial B, there were 28 trees per plot row (with a total of 84 in each experimental plot). Four blocks of each treatment were randomized at each site. Six fruit clusters of each irrigation treatment were collected at 140 dpa, near commercial harvest.

The ID number of the trees sampled from USDA Pistachio Germplasm Collection located at Wolfskill Experimental Orchards (Winters, CA, USA) are provided in Supplemental Table 1. Fruits were collected and scored from 2021-2024 growing seasons. Three - 6 fruit clusters throughout the tree were bagged and manually pollinated and 2 - 6 clusters were collected at 147-154 dpa.

### Sample fixation and sectioning

Whole fruits were imaged and excised into either 5 mm x 5 mm segments or 5 mm rings in the transverse plane, imaged again in the transverse plane, and chemically fixed. The samples were vacuum infiltrated (Labcanco, model 5530000, USA) at a pressure of -90 to -100 KPa for 1.5 hrs at RT in fixative solution containing 4% (W/V) paraformaldehyde (PFA) (Thermo Scientific, formerly Acros Organics #M26863, USA) in phosphate buffer saline (PBS) and 0.1% Tween-20. Fixed samples were washed for 10 min in double deionized water prior to embedding. A vibratome (Leica Biosystems Nussloch GmbH, VT1000S, Germany) was used to section tissue to 100 μm as previously described (Zhang et al., 2021). For sectioning of both fresh and fixed samples, the samples were trimmed down to 3-5 mm around the region of interest and embedded in 5% (W/V) agarose (Sigma-Aldrich, #A9539, USA) in distilled and deionized water.

### Hull firmness measurement

Hull Firmness was measured on a Texture Analyzer (TA.XT Plus, Texture Technologies Corp. Hamilton, MA) with TA-52 probe (2 mm diameter), using the software Exponent (Version 6.1.16.0; Texture Technologies Corp.). Each fruit was split into two halves along the suture line so the samples would lie flat and placed on the platform of texture analyzer. The probe was set at the speed of 1 mm s^-1^ to penetrate the hull, shell, and the kernel, and the first peak, indicating the force necessary to penetrate the hull, was recorded as hull firmness as described by Adaskaveg et al <u>(Adaskaveg et al., 2025)</u>.

### Cellular staining

Samples were stained with undiluted Calcofluor White containing Evan’s blue (Sigma-Aldrich, #18909, USA) for 5 min at RT and rinsed three times with distilled and deionized water before imaging. Toluidine blue (Sigma-Aldrich, #T-3260, USA) staining was performed as described by Mitra & Loque 2014 (Pradhan Mitra & Loqué, 2014). Briefly, sections were incubated in 0.02% (W/V, in distilled and deionized water) toluidine blue solution for 2 to 5 min at RT then rinsed three times with distilled and deionized water before imaging. Sudan IV (Matheson Coleman & Bell, #B 417, USA) staining was performed by incubating sections for 5 min in 0.09% Sudan IV in 1:1 ethanol:glycerol at RT as described by Ruzin et al (Ruzin, 1999). Fluorol Yellow 088 staining was performed as described previously (Zhang et al., 2021). Briefly, the sections are incubated in 0.1 mg mL^-1^ Fluorol Yellow 088 (Santa Cruz Biotechnology CAS-81-37-8, USA) in lactic acid at 70 °C for 1 h in the dark, rinsed twice with distilled and deionized water, before imaging mounted in 50% glycerol (V/V, in distilled and deionized water).

### Immunofluorescence

100 μm sections of fixed tissue were stained with antibodies based on protocol modified from Gerttula & Groover (Gerttula & Groover, 2017). Briefly, the sections were rinsed in distilled and deionized water for at least 15 min before washing three times with PBS 0.2% Tween-20 and blocking in blocking buffer (2% fish gelatin (Fisher Scientific, #50-259-35, USA), 1% bovine serum albumin (Chem-Impex, #00040, USA), 0.2% Tween-20 in PBS) for 1 h at RT. Pectin staining was performed using JIM5 antibody (PlantProbes, University of Leeds, Leeds, UK) against low methylesterified homogalacturonan at 1:10 dilution in blocking buffer, and JIM7 antibody (PlantProbes) against highly methylesterified homogalacturonan at 1:8 dilution in blocking buffer. Staining with primary antibody was performed overnight at 4 °C in a humid chamber before washing three times with PBS 0.2% Tween-20 and blocking for 1 h at RT. The anti-rat AlexaFluor 488 (Invitrogen, #ab150157, USA) was used as a secondary antibody for both JIM5 and JIM7 at 1:50 dilution in the blocking buffer. The secondary antibody staining was performed overnight at 4 °C in the dark before washing three times in PBS 0.2% Tween-20, mounting in the same PBS-Tween mixture, and imaging.

### Microscopy

All bright field micrographs (toluidine blue, Evan’s blue, Sudan IV) were captured using Olympus BH-2 microscope with Jenoptik D-07739 microscope capture camera and the ProgRes CapturePro 2.10.0.1 software (Jenoptik AG, Germany) and SPlan 4x 0.13, 10x 0.3, and 40x 0.7 dry objectives. Fluorescence microscopy images were acquired using the confocal Laser Scanning Microscopy 980 system (Carl Zeiss, Germany). A 408 nm laser (2% power) was used to capture the Calcofluor White signal and a 488 nm laser (8% power) was used for goat anti-rat secondary antibody, Alexa Fluor™ 488 for JIM5 and JIM7. Emissions were collected over a wavelength range of 410 to 488 nm for Calcofluor White and 494 to 582 nm for the Alexa Fluor^TM^ 488 secondary antibody of JIM5 and JIM7. All fluorescence images were collected using either the Plan-Apochromat 10x 0.45 M27 objective or the Plan-Apochromat 20x 0.8 M27 objective. The pinhole was set to 1.88 AU for Calcofluor White, 11.16 AU for JIM5 and JIM7. Zen 2011 SP3 (Zeiss) software was used for imaging and Zen Lite (ZEN 3.2) for image export, with JIM5 and JIM7 exported at display threshold of 15,000. Autofluorescence micrographs were captured on three channels. The blue channel was excited with a 405 nm laser at 7% power, 10.59 AU pinhole, and emission collected from 410 - 483 nm. The green channel was excited with a 488 nm laser at 10% power, 4.4 AU pinhole, and emission collected from 494 - 582 nm. The magenta channel was excited with a 561 nm laser at 10% power, 4.72 AU pinhole, and emission collected from 573 - 719 nm.

### Quantification of anatomical traits

Hull thickness and cell size were measured manually using the “measure” function in FIJI (Fiji Is Just) Image J (1.53f51) (Schneider et al., 2012). Images and micrographs of fruits sectioned in the transverse plane, at the median of the fruit between the apex and base of the fruit, were used. The dorsal site is defined at ∼ 2 mm region around the suture ridge on the pistachio with the longer curvature, ventral site is ∼ 2mm region around the suture ridge on the pistachio with the shorter curvature, and the nonsuture region is defined as the 4 mm region located 90° away from both dorsal and ventral suture. The polygon tool was used to select the cells in the micrograph for cell cross section area measurements. The line tool was used to measure the thickness of the tissue of interest, by drawing a line at the start and end of the tissue layer, perpendicular to the epidermis.

### RNAseq

’Golden Hills’ and ’Lost Hills’ fruits were harvested from the trees tagged in the 2022 commercial orchard between 11 am and 1 pm at 91 and 133 dpa and transported on ice back to the lab, where the hull was removed and snap-frozen in liquid nitrogen.

At 91 dpa, all the fruits had intact hulls. At 133 dpa the fruits were separated by hull phenotype into three categories: 1) cracked, 2) intact, and 3) tattered. Four fruits from each tree were pooled for each biological replicate and four biological replicates, representing four trees, were analyzed per category at each time point. Due to the frequency of hull split events, only ’Golden Hills’ was used for tattered category, and only ’Lost Hills’ for cracked category. Total RNA was extracted from hull tissue using RNeasy Plant Mini Kit (Qiagen, #74904, USA) and sent to Novogene Corporation Inc (Sacramento, CA, USA) for RNA-seq library preparation, sequencing, and bioinformatics analysis. The analysis utilized the initial version of *P*. *vera* ‘Kerman’ genome assembly and annotation, and was then updated to incorporate the most recent gene annotations following publication of the ‘Kerman’ genome and annotation (Adaskaveg et al., 2025). RNAseq quality statistical calculations, correlation plots, Differential gene expression (DGE) analyses, Gene Ontology analyses, and KEGG pathway analyses were performed by Novogene. Briefly, DEG was performed using DESeq2 (Love et al., 2014) with |log2(FoldChange)| >= 1 & adjusted P value <= 0.05 set as the threshold. GO and KEGG were both performed using the clusterProfiler (Yu et al., 2012). For GO, and KEGG, GO terms and KEGG pathways with adjusted P value < 0.05 were considered significant enrichment. The top 30 GO terms were visualized as dot blots and the top 20 KEGG pathways were visualized as bar plots for each analysis.

Read counts of candidate genes in Fragments Per Kilobase Million (FPKM) identified in DGE were then visualized using the ggplot2 package (Wickham 2016) in R as heatmap.

### qRT-PCR

Samples for qRT-PCR analyses were harvested and processed for RNA following the same protocol as described in the RNAseq section above. cDNA synthesis was performed using SuperScript III First-Strand Synthesis System (ThermoFisher Scientific, #18080051, USA) following the manufacturer’s protocol, with RNase H digestion as a final step to eliminate residual RNA. Primers are designed using Primer-BLAST (National Center for Biotechnology Innovation) using published *P. vera* sequences in the RefSeq mRNA database (National Center for Biotechnology Innovation) for genes of interest, with one primer bridging an exon-exon junction to avoid detection of residual genomic DNA. The primers used are listed in Sup. Table 2. qPCR is performed using Power SYBR Green PCR Master Mix (ThermoFisher Scientific, #4368577) using the standard two-step amplification protocol for 10 µL reactions, as per instruction from the manufacturer, on the CFX96 Real-Time System with C1000 Touch Thermal Cycler (Bio-Rad, #1845096, USA).

Since we detected amino acid and nucleotide sugar metabolism, motor protein, and MAPK signaling pathways as significantly different between hulls of different categories in our KEGG analysis (Sup. Fig. 1), we questioned whether housekeeping genes commonly used as reference genes will still show stable expression between our test samples. We analyzed all the read counts of actin, *EF1*α, and *GAPDH* gene families, which are routinely used as reference genes for qRT-PCR, and found that while actin and *GAPDH* genes displayed stable expression during late stage development and hull split, *EF1*α showed significant variation between time points and between cracked and tattered hull (Sup. Fig. 1). Thus, qRT-PCR were normalized using *ACT7* (PvKer.03.g088750), chosen due to its high read count and stable expression.

### Cell adhesion test

Cell-cell adhesion of the pistachio hull was modified based on protocol from Segonne et al (Segonne et al., 2014). Briefly, 5 x 5 mm of hull tissue was removed from the nonsuture site at the median of the fruit, halfway between the tip and the base, submerged in 2 mL of deionized water and vortexed (Fisher Scientific, Model G550) at maximum speed for 2 minutes. A 30 μL aliquot of the cell suspension was then mounted on a glass slide and the number of non-adherent cells released from the tissue counted.

### Weather data collection

The daily weather conditions for the 91 dpa to 133 dpa period in 2022-2024 were collected from the California Irrigation Management Information System (CIMIS) weather station #139 (California Department of Water Resources 2023), located at the site of the USDA Germplasm Collection in Wolfskill Experimental Orchard (Winters, CA, USA). Growing Degree Days (GDD) data for 2021 and 2022 commercial orchards used in this study are as published by Zhang et al (Zhang et al., 2023).

### Water exposure and solar radiation assays

<u>Hull integrity after acute water exposure</u> was tested by spraying the fruit cluster of seven trees from the USDA Pistachio Germplasm Collection at Wolfskill Experimental Orchard with deionized water at 91, 112 and 133 dpa during the 2024 growing season, to correspond to the time points for transcriptome analysis, with an additional time point added at 112 dpa to increase the frequency of water exposure. The fruits were sprayed until thoroughly wet, to simulate rain conditions, before they were allowed to air dry. One cluster was sprayed per tree while another cluster close by was used as control. The sprayed clusters were harvested at 142 dpa and scored for hull phenotype. <u>Hull integrity after chronic water exposure</u> was tested by covering one fruit cluster per tree of eight ‘Golden Hills’ trees during the 2024 growing season in 10 none-dripping wet paper towels and then enclosed in glassine paper bags (ULINE, Pleasant Prairie, WI, S-12472, 4.5 Liter) to mimic a prolonged high humidity condition. Unbagged clusters were used as control. The towels are moistened with 50 mL deionized water at 2 - 4 biweekly time points at 91, 105, 119, 133 dpa before harvest at 141 dpa and phenotypically analyzed for hull degradation integrity.

### Statistical analysis & figure assembly

R x 64 version 4.0.3 (R Core Team, 2017) in R studio (RStudio, PBC), version 1.3.1093 was used for statistical analyses. The basic ANOVA, T-test functions and the emmeans (version 1.6.3) and multcomp (version 1.4-17) packages were used for statistical analysis. Final graphs were generated using Microsoft Excel Office 365 (Microsoft, Redmond, WA, USA) with the exception of heatmap, which was generated using the ggplot2 package (Wickham 2016) in R.

Photographs and micrographs were formatted in GIMP version 2.10.38 graphics editor and FIJI (Fiji Is Just) Image J (1.53f51) (Schneider et al., 2012). All figures were assembled and edited in Inkscape version 1.0.1 (3bc2e813f5, 2020-09-07, open-source software).

## Results

### Two main categories of hull split in pistachio cultivars

To understand late-stage pistachio hull development and the mechanisms underlying hull split, we first characterized and categorized the different hull split phenotypes that occur during late-stage fruit development during the 2022 growing season. We selected ‘Kerman’ as the industry standard nut cultivar, with a recently published high quality genome (Adaskaveg et al., 2025), and compared it with ‘Golden Hills’ (Kallsen et al., 2009) and ‘Lost Hills’ (Kallsen et al., 2014), two upcoming cultivars with earlier harvest dates and higher shell split rates. We set our comparison at 133 dpa at the end of August, coinciding with ‘Golden Hills’ commercial harvest (∼ 2000 GDD). Our analysis revealed four main categories of hull phenotypes: 1) intact, 2) cracked, where the exo-mesocarp was completely split along the interior-exterior axis so that the endocarp (shell) was exposed, and 3) tattered, with superficial split, largely at the exocarp layer, while the shell and kernel remained protected and 4) combination of both cracking and tattering (Fig. 1A). The split occurred between 91 and 133 dpa, and was consistent in appearance between genotypes (Sup. Fig. 2A, B) The frequency of occurrence for each category appeared cultivar specific, with ’Golden Hills’ showing the highest percentage of tattering at 16%, a cracking rate of 2% and intact hull rate of 79% (Fig. 1B). In contrast, ’Lost Hills’ showed a reverse order with 16% cracking, 1% tattering and 78% intact (Fig. 1B). ‘Golden Hills’ and ‘Lost Hills’ displayed comparable frequencies of fruits with combined tattering and cracking (Fig. 1B). In contrast to these cultivars, ’Kerman’ showed a significant phenotypic difference with 100% hull integrity at 133 dpa, without any type of hull split (Fig. 1b, P < 0.01 Fisher’s Exact Test). A second time point was performed at 142 dpa (∼2100 GDD), allowing further characterization of late stage ‘Kerman’ hull development due to its later harvest date. The frequency of hull split increased from 133 to 142 dpa as expected, since the ripening processes continued. However, the cultivars maintained the differences in hull split categories with the same trend (Fig. 1C, P < 0.01, Fisher’s Exact Test). ’Golden Hills’ showed a higher rate of tattering (32%) than cracking (18%), while ’Lost Hills’ showed a higher rate of cracking (28%) than tattering (15%) (Fig. 1C). Notably, ’Kerman’ began to show hull cracking (8%), but not tattering at 142 dpa (Fig. 1C).

**Fig. 1.**
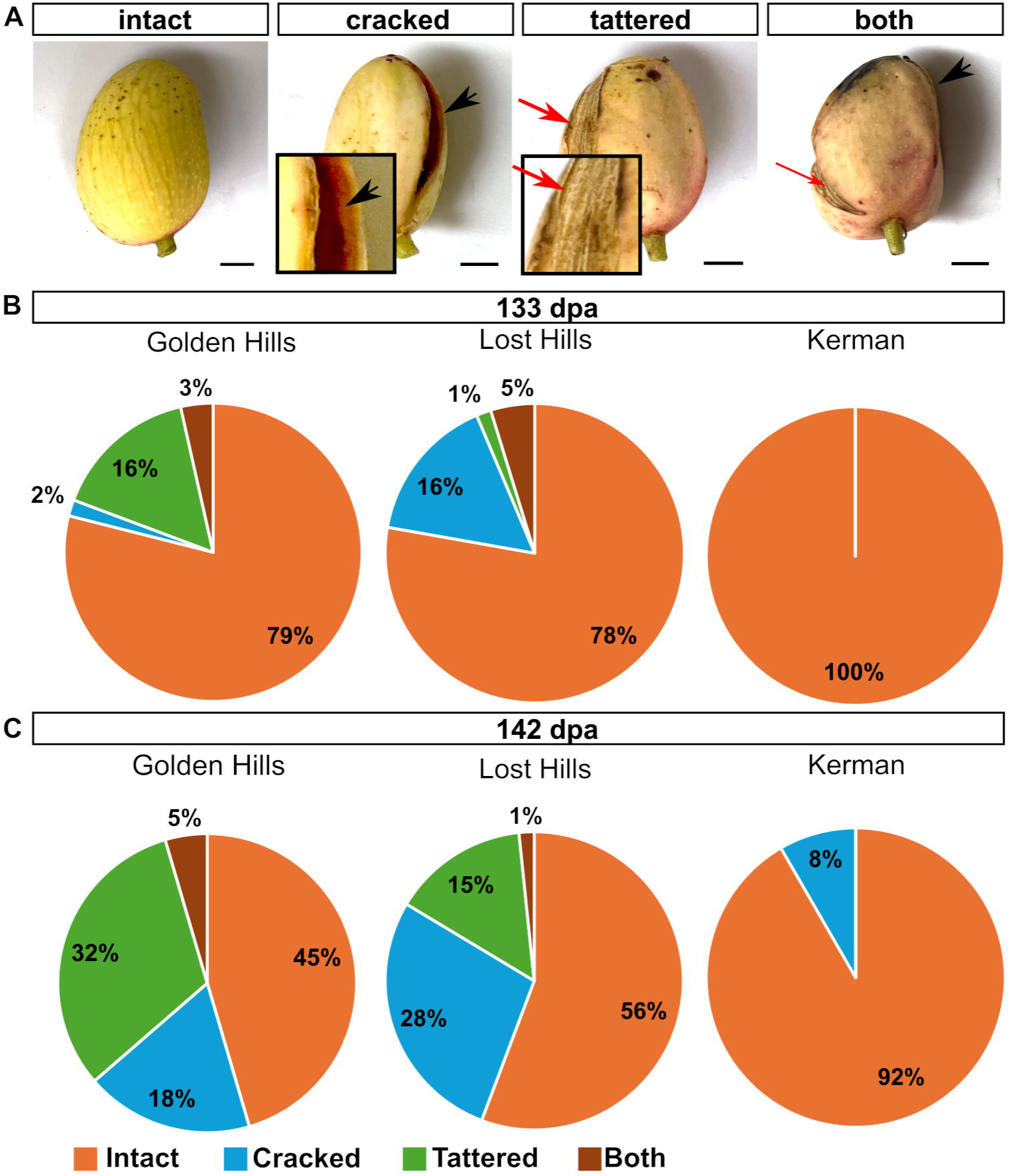
Phenotype of pistachio hull split during late-stage fruit development. (A) Representative images of pistachio fruits demonstrating the different categories of hull integrity at 133 days post anthesis (dpa). The site of hull cracking (black arrow) indicates a deep split while hull tattering (red arrow) is superficial. Insets show representative areas of split. ‘Golden Hills’ fruits are used for all except cracked hull, which is ‘Lost Hills’. Scale bars = 5 mm. (B) Frequency of occurrence for the different hull integrity categories observed in 2022 at 133 dpa (B) and 142 dpa (C). N = 24-63 fruits per genotype per time point. P < 0.01 for each time point. Fisher’s Exact Test.

Due to the late maturation of ‘Kerman’ and to account for the difference in orchard and weather between the years, we included a further sampling date at 154 dpa (∼3000 GDD) for ‘Kerman’ from a different growing season to allow for a saturation point of maturity, and to test if ’Kerman’ will display a similar increase in cracking. ‘Kerman’ showed 100% integrity at 133 dpa and a hull split of 38% of fruits showing primarily tattered hulls at 154 dpa (Sup. Fig. 2C).

### The pistachio exocarp contains multi-layered hypodermal and filler parenchyma tissue

Given the importance of the pistachio hull as the tissue layers that encapsulate and protect the endocarp and the seed (Doster & Michailides, 1995, 1999), we aimed to create a clear anatomical definition of the hull and its split categories. Thus, we examined the hull cell type composition and established its anatomical structure to better understand the cellular mechanisms behind each split type. Cell staining and anatomical analysis showed that the pistachio hull is composed of the exocarp and the mesocarp with many cell types (Fig. 2). The cuticle, epidermis, and hypodermal parenchyma form the skin, or exocarp, in an organization pattern similar to other fruits (Santos et al., 2023). We defined the mesocarp in pistachio to include all the cells between the exocarp and the endocarp (shell). Hence, the hull consists of all the cell layers encompassing both the exocarp and the mesocarp, namely the exocarp hypodermal parenchyma and the mesocarp filler parenchyma (Fig. 2A). The hypodermal parenchyma (HP) in the exocarp tends to be smaller, more flattened, and can contain 3-7 cell layers (Fig2A-ii, iv, blue cell outline). The filler parenchyma (FP) in the mesocarp, located interior to the HP, contains 18-26 cell layers that are larger and more isodiametric (Fig2A-ii, iv, red cell outline). In between the HP and FP is the transition zone, containing 3-5 parenchyma cell layers with phenotype that is intermediate between HP and FP (Fig 2A). The transition zone tends to be the site of deposition for phenolics and pigments with high autofluorescence properties and the site where cell senescence initiates during late-stage hull development (Fig. 2A-iii, iv). Thus anatomically defined, hull cracking involves the splitting of the entire exo-mesocarp (Fig. 2B), whereas hull tattering corresponds to the breakdown and separation of the HP from FP along the transition zone (white dashed line, Fig. 2B), resulting in the separation of the exocarp from the mesocarp.

**Fig. 2.**
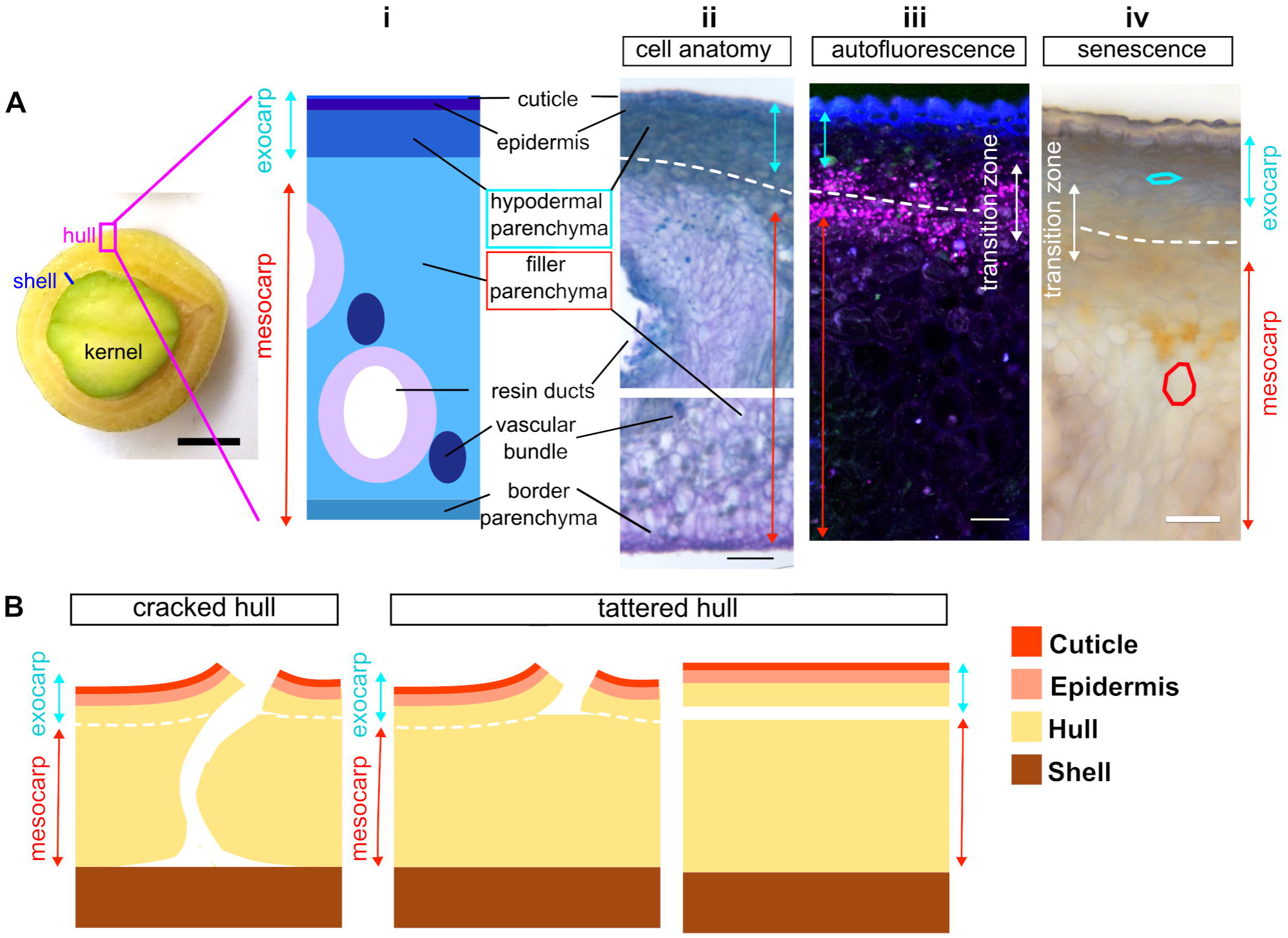
Anatomy of pistachio hull during the late-stage development. (A) Diagram and transverse sections from nonsuture region (magenta box) depicting the anatomical features of the hull of 133 days post anthesis pistachio (i). Scale bar = 5 mm. Cell anatomy is visualized by toluidine blue staining (ii), scale bar = 100 µm, and cell autofluorescence (iii), scale bar = 50 µm. Cell senescence is visualized with Evans blue staining (iv), scale bar = 50 µm. The exocarp (skin, cyan) contain the cuticle, epidermis, hypodermal parenchyma , while the mesocarp (red) contains resin ducts, vascular bundles and their associated accessory tissues as well as filler and border parenchyma. The transition zone is indicated in white, while the dashed line indicates one potential line of exocarp mesocarp separation during hull tattering. (B) Schematic representation of the cracked and tattered hull events, with the transition zone indicated with white dashed line.

### RNAseq identifies cell wall modification genes associated with pistachio late-stage hull development

To better understand the interaction between hull integrity and late-stage development, we performed RNAseq analysis. We used ‘Golden Hills’ and ‘Lost Hills’ harvested in 2022, which showed cultivar-specific differences in hull split categories (Fig. 1). We selected 91 dpa, when all hulls were intact, and compared it against 133 dpa, when hull split had occurred, to examine differences in development between intact and split hulls. We focused the analysis of hull cracking on ’Lost Hills’ and the analysis of hull tattering on ’Golden Hills’ because they displayed high frequency of these phenotypes (Fig. 1B, C). Statistical qualities for the RNAseq are reported in Sup. Table 3. On average we were able to obtain 42,751,562 raw reads per sample, of which 86.8% were mappable to the ‘Kerman’ reference genome. Pairwise DGE analysis between genotype, dpa, and phenotype showed that the greatest number of differentially expressed genes was detected between 91 and 133 dpa tattered ’Golden Hills’ hull samples (6128 genes, Sup. Fig. 3A, red arrow), while the smallest number of differentially expressed genes was detected between ’Golden Hills’ and ’Lost Hills’ hull at 91 dpa (699 genes, Sup. Fig. 3A, black arrow). In contrast to the large number of genes that are differentially expressed between 91 and 133 dpa tattered ‘Golden Hills’ (red arrow), fewer differentially expressed genes were found in analyses comparing 133 dpa intact hulls of ‘Golden Hills’ and ‘Lost Hills’ hull to their corresponding 91 dpa samples (4658 and 3179 genes, Sup. Fig. 3A, cyan arrows). Pearson’s correlation analysis likewise showed that the genotypes are similar at 91 dpa, while the greatest difference in transcriptome is between 91 dpa ’Golden Hills’ and 133 dpa ’Golden Hills’ tattered hull (Sup. Fig. 3B).

We then looked at the top 50 genes, sorted either by adjusted P value or by fold change, from the pairwise DGE analysis across time points, genotypes, and phenotypes. The top ten candidates are listed in Sup. Tables 4-6. Several genes among the top ten by P value candidates were differentially expressed across multiple analyses, including two known to be involved in cell wall remodeling (Fig 3A). Snakin-1 (*SNAK1,* PvKer.10.g252770) showed increased expression from 91 to 133 dpa in intact ‘Golden Hills’ (log2 fold increase 6.03 adjusted P = 3.21E-139) and from 91 to 133 dpa in tattered ‘Golden Hills’ (log 2 fold increase 7.28, adjusted P = 1.25E-275) (Fig 3A). Expansin 4 (*EXPA4,* PvKer.03.g089030) showed a log2 fold increase of 4.1 between intact and tattered 133 dpa ‘Golden Hills’ (adjusted P = 3.35E-66), and log 2 fold increase of 3.51 from intact and cracked (adjusted P = 1.21E-19) at 133 dpa ‘Lost Hills’ (Fig. 3A).

**Fig. 3.**
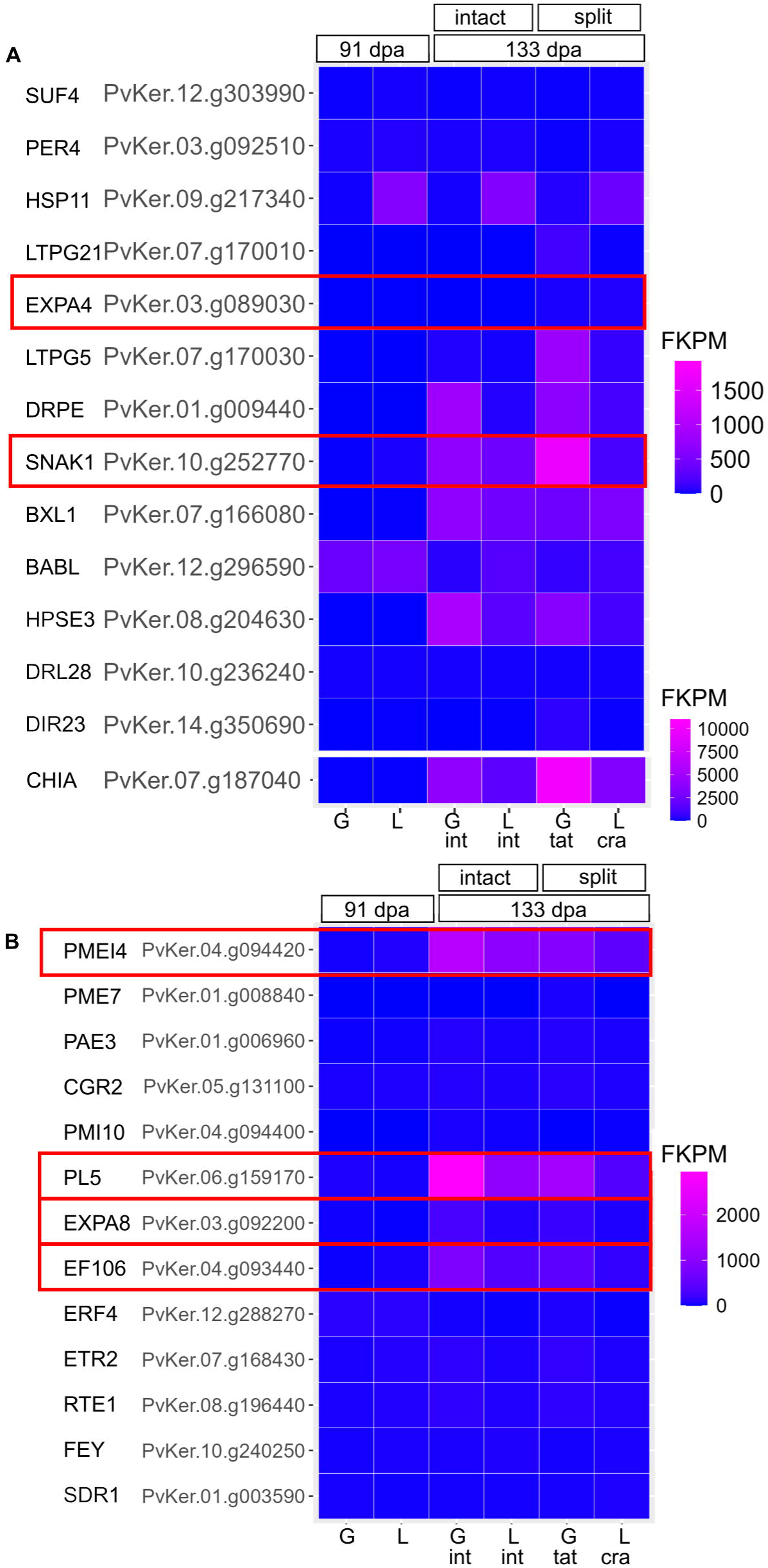
Expression of top candidate genes for pistachio fruit during late-stage development. (A) Heatmap of the Fragments Per Kilobase per Million mapped reads (FKPM) of top candidate genes identified in multiple differential gene expression analyses in RNAseq. ‘Golden Hills’ = G, ‘Lost Hills’ = L, intact hull = int, tattered hull = tat, cracked hull = cra. CHIA (acidic endochitinase) is re-plotted on a separate scale to avoid drowning the signal of the other candidate genes. (B) Heatmap of the FPKM of additional genes of interest that showed differential expression for cell wall modification and hormonal responses. Each square represents the average of four biological replicates.

Since all hulls were intact at 91 dpa, we reasoned that fruit ripening processes may be associated with hull split. Therefore, we also examined the genes involved in pectin component of the cell wall, ethylene biosynthesis, and additional genes in expansin family in our DGE results, as they are implicated in fruit ripening (Brummell et al., 1999; Civello et al., 1999; Cosgrove, 2000; Forlani et al., 2019; Jiang, Lopez, Jeon, de Freitas, et al., 2019; Jiang, Lopez, Jeon, De Freitas, et al., 2019; Santiago-Doménech et al., 2008; Wang et al., 2018, 2019). We observed cultivar-specific expression up-regulation from 91 to 133 dpa for Pectin Lyase 5 (*PL5,* PvKer.06.g159170, log 2 fold change 5.37, adjusted P value = 5.97E-106), Pectin MEthylesterase Inhibitor 4 (*PMEI4,* PvKer.04.g094420, log 2 fold change 5.04, adjusted P value =1.1E-49) and Ethylene responsive transcription Factor 106 (*EF106,* PvKer.04.g093440, log 2 fold change 4.65, adjusted P value = 1.27E-47) (Fig. 3B), and that the upregulation is most significant comparing 91 to 133 dpa intact ‘Golden Hills’. Expansin 8 (*EXPA8,* PvKer.03.g092200) also showed up regulation in this analyses (log 2 fold change = 2.58, adjusted P value = 1.77E-12), and it is among the top 50 genes by P value in our analysis of cracked versus tattered fruit (log 2 fold change = 1.77 higher in tattered, adjusted P value = 1.29E-38). Interestingly, we found two genes that may be involved in pectin biosynthetic genes that were up regulated from 91 to 133 dpa, though they were not among the top ten in the DGE analyses. Galacturonosyltransferase 3 (GAUT3, PvKer.14.g345950) was up-regulated by two-fold (adjusted P value = 1.27E-17) from 91 to 133 dpa, suggesting that multiple pectin modification processes, including synthesis of new pectin, occur during late-stage development.

Next, we validated the expression of *PL5,* for pectin lyase’s well-established role in ripening, *SNAK1,* which is associated with both plant defense and cell wall remodeling response (Almasia et al., 2008; Nahirñak et al., 2012), and *EXPA8,* as the candidate to test for differences between cracked and tattered fruits. We used qRT-PCR to test ‘Golden Hills samples’. Similar to RNAseq, there was a significant upregulation of *PL5* at 133 dpa compared to 91 dpa, when intact hull had higher *PL5* expression than tattered hull (Sup. Fig. 4). *SNAK1* had a threshold significance of P = 0.055, with higher expression for the tattered fruits, corroborating with the trend we observed in RNAseq (Sup. Fig. 4). However, changes in *EXPA8* were below our qRT-PCR detection threshold (Sup. Fig. 4).

We used gene ontology (GO) and KEGG analysis to identify pathways associated with hull phenotypic traits. There was a greater difference in hydrolases acting on for hydrolyzing O-glycosyl compounds and glycosyl bonds (GO:0004553, GO:0016798) when tattered hulls were compared against intact or cracked hulls at 133 dpa in GO (Sup. Fig. 5A, C, E). Interestingly, the phenylpropanoid and flavonoid pathways showed the most pronounced differences when the intact hull was compared to all other phenotypes (Sup. Fig. 5B, D), potentially due to stress caused by ruptured tissue. In contrast, plant hormonal and MAPK signaling pathways were the most altered when cracked hull was compared to tattered hull (Sup. Fig. 5F), suggesting that cell cycle / senescence signaling may be different between cracked and tattered hulls.

### Fruit softening is associated with hull tattering but not cracking

Since we detected significant differences in cell wall remodeling genes during late-stage development of pistachio hull, we next investigated the hull texture of fruits obtained during the 2022 growing season to determine if there was a cultivar-specific difference in the texture. We observed that hulls softened significantly from 91 to 133 dpa in all genotypes, and it was most dramatic in ’Golden Hills’, with nearly a five-fold decrease in penetration force from 91 dpa to 133 dpa (Fig. 4A), consistent with our transcriptome results.. In contrast, ’Lost Hills’ required around twice as much force to penetrate at 133 dpa compared to ’Golden Hills’ (Fig. 4A). To distinguish the impact of genotypes on the phenotype-specific differences, we separated the 133 dpa fruits by their hull split phenotypes. Even though this significantly reduced the sampling size, we were still able to detect that regardless of the cultivar, intact hull always required the most force for hull penetration, while tattered hull required the least force, suggesting that hull softening is associated with separation of the exocarp from the mesocarp. Cracked hulls were more similar in texture to intact hulls, while hulls that showed both tattering and cracking showed a texture that is more similar to tattered hulls (Fig. 4B-D).

**Fig. 4.**
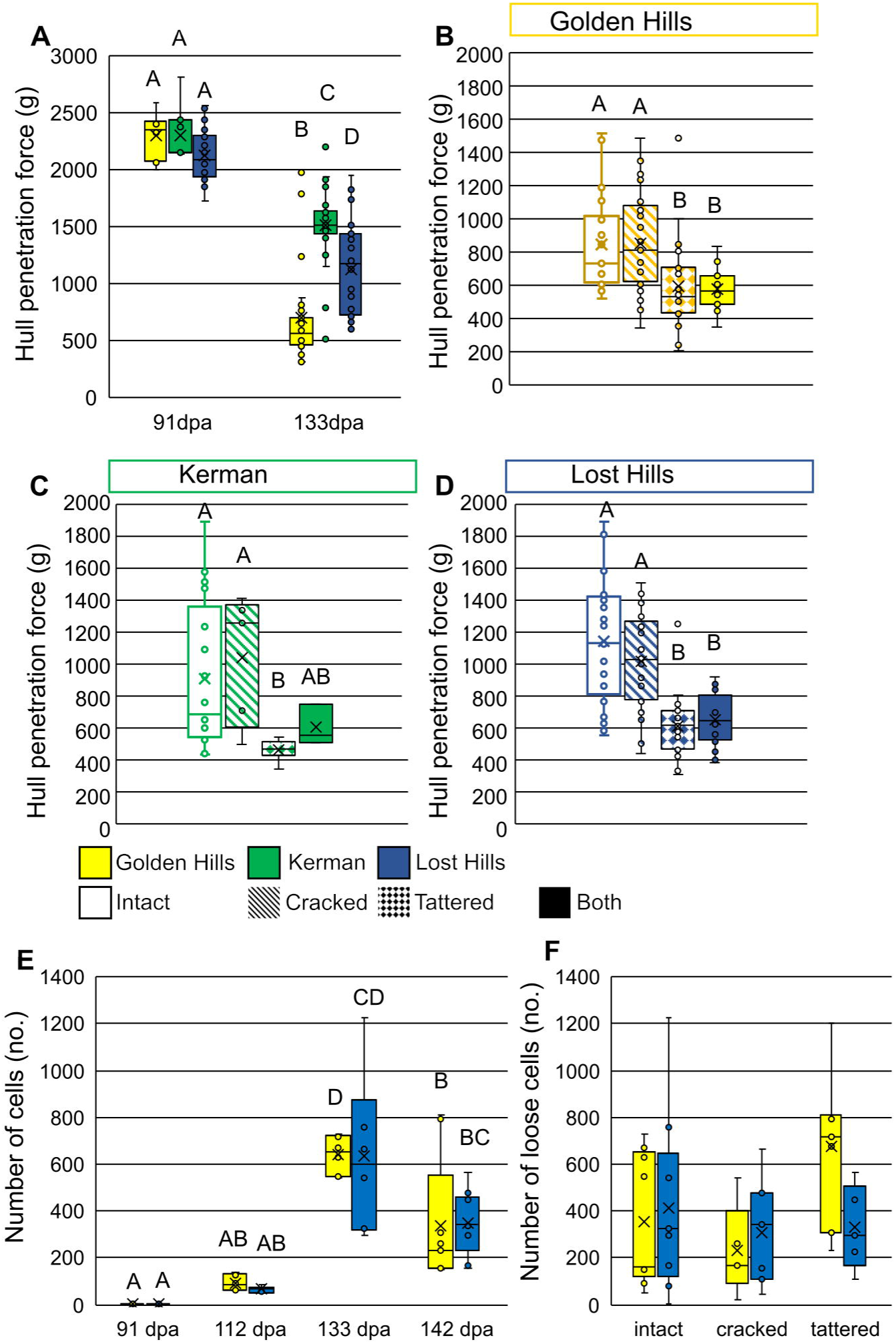
Changes in hull texture and cell-cell adhesion during hull late-stage development. (A) Changes in hull texture during ripening measured using hull penetration force for ‘Golden Hills’ (GH), ‘Kerman’ (K), and ‘Lost Hills’ (LH). N = 10-36 fruits from 6 trees. P < 0.01 for time points, genotype, two-way ANOVA. (B-D) Textural differences between different hull phenotypes for cultivars GH (B), K (C), and LH (D). N = 44-91 fruits per cultivar. One-way ANOVA. P < 0.05. (E) Number of non-adherent cells released in cell-cell adhesion assay during hull ripening stage in 2022. N = 6-9 fruits per cultivar per time point. Two-way ANOVA. P < 0.01 between time points, NS for cultivar and interaction. (F) Number of non-adherent cells released in cell-cell adhesion assay for 133 - 142 dpa ’Golden Hills’ and ’Lost Hills’ fruits with different hull integrity, harvested in 2022. P is NS for all. Two-way ANOVA. N = 7-11 fruits. All samples are from the 2022 growing season. Different letters indicate Least Square Means post-hoc analysis significant with α=0.05

Fruit softening during ripening and pectin modification is often associated with a decrease in cell-cell adhesion (Harker & Hallett, 1992; Lahaye et al., 2021; Segonne et al., 2014; Su et al., 2024). Therefore, to test whether cell-cell adhesion also decreases during late stage hull development and whether it is phenotype-specific, we measured the number of non-adherent cells, or cells that are separated from the main hull tissue, using an established cell-cell adhesion assay (Segonne et al., 2014). A nearly 600-fold increase was detected in the number of non-adherent cells for all hull phenotype categories from 91 dpa to 133 dpa when all phenotype categories were pooled (Fig. 4E), however, there was no genotype-specific difference (Fig. 4E).

When 133-142 dpa fruits were separated by their genotype and phenotype, tattered ’Golden Hills’ hull had a higher number of non-adherent cells, albeit this was not statistically significant (Fig. 4F).

### Late-stage development and higher split rate is associated with increased pectin detection signal

Our data suggested significant pectin remodeling occurred during late-stage hull development. To better understand the type of pectin modification involved in pistachio hull split, we investigated the pectin changes by immunofluorescence with pectin-specific antibodies on transverse sections of ’Golden Hills’ and ’Lost Hills’ fruits at 91 dpa, when all hulls were intact, and 133 dpa, when hull split has occurred. Employing the monoclonal antibody, JIM5, which detects low methylesterified pectin (Clausen et al., 2003; Knox et al., 1990), we found a fluorescence signal increase from 91 to 133 dpa, especially at the cell wall junctions and in the transition zone between the HP and FP cell layers (Fig. 5A, white arrow). Similarly, an increase in the JIM7 immunofluorescence signal, which detects high methylesterified pectin, was observed for both genotypes (Fig. 5B) (Clausen et al., 2003; Knox et al., 1990). Quantification of fluorescence showed that while both JIM5 and JIM7 signal intensity increased from 91 to 133 dpa, cell type responses differed. For JIM5, the FP signal increased significantly. (Fig. 5C) For JIM7 the HP fluorescence increased significantly, with JIM7 also showing a cultivar-specific difference in total hull signal at 133 dpa (Fig. 5C). When we compared hull phenotypes, JIM5 showed genotype specific differences between ‘Golden Hills’ and ‘Lost Hills’ intact and cracked fruits, and there was a slight trend of increased fluorescence signal in ‘Golden Hills’ split hull compared to intact hull, albeit not statistically significant. (Sup. Fig. 6A) JIM7 staining showed similar levels in intact fruits of both genotypes, and differences were observed between hull phenotype categories when hull split occurred, with a nearly fourfold change in cracked hull compared to intact hull in ‘Lost Hills’ and nearly fourfold change in tattered hull compared to intact hull in ‘Golden Hills’ (Sup. Fig. 6B)

**Fig. 5.**
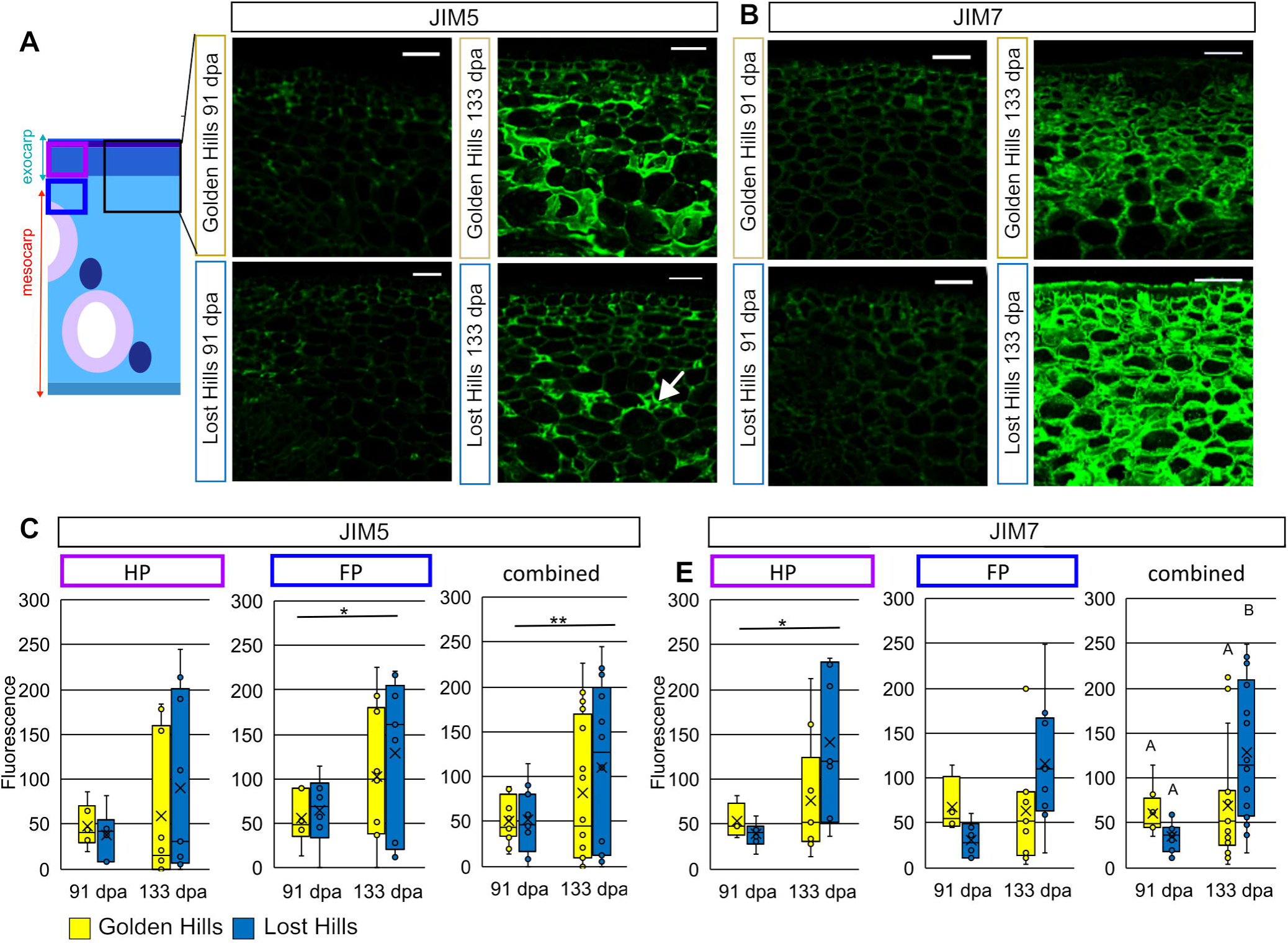
Spatiotemporal analysis of pectin modification. (A) Hull sections stained with JIM5 for low-methylesterified pectin at 91 and 133 days post anthesis (dpa). White arrow indicates signal at cell-cell junction. Diagram indicates region of hull sampled. (B) Imaging of hull sections stained with JIM7 for high-methylesterified pectin. Scale bars = 50 µm. (C) Quantification of average JIM5 and JIM7 fluorescence intensity for hypodermal parenchyma (HP), filler parenchyma (FP) , and whole micrograph from samples harvested in 2022. All phenotype categories are included for (B) and (C). * P < 0.05, ** P < 0.01, Two-way ANOVA. Different letters indicate Least Square Means post-hoc analysis significant with α = 0.05, no letter indicates NS in post-hoc analysis. N = 6-10 sections quantified from fruits of 5-6 trees.

We further examined the association of hull pectin in different growing seasons and observed that during 2021, the growing season in which a high percentage of fruits with intact hulls occurred (Sup. Fig. 2C), the average fluorescence intensity appeared unchanged from 91 dpa to 133 dpa (Sup. Fig. 6C, D). Thus, environmental differences between growing seasons could contribute to differences in pectin modification during late-stage hull development, and affect hull split rate.

### Effects of expansion, hull thickness, and cuticle on hull split

Given that we identified expansin and pectin remodeling genes in our RNAseq experiment along with the changes in the levels of pectin (Fig. 3, 5), we wanted to investigate whether cell and tissue expansion contribute to hull split phenotypes.

Since the pistachio hull represents a structurally complex tissue with an expanded number of HP layers relative to the simpler organization observed in berries (Fig. 2) (Bargel et al., 2004; B.-M. Chang & Keller, 2021; Konishi et al., 2022; Wang et al., 2019), we hypothesized that cell type-specific changes in size may contribute to the internal expansion force driving hull split. We compared the cell area of FP against HP in 91 and 131 dpa for ‘Golden Hills’ and ‘Lost Hills’ sampled from the 2022 growing season. The FP cell size nearly doubled from 91 to 133 dpa, while the HP size remained constant (Fig. 6). A similar trend was observed in 2021, the growing season with high integrity. During this high integrity season, FP increased in size, albeit not statistically significant, while HP remained constant in size (Sup. Fig. 8). In summary, we found that there are cell-type-specific differences in cell size expansion during late-stage fruit development, where FP cell size in the mesocarp increased while HP in the exocarp did not.

**Fig. 6.**
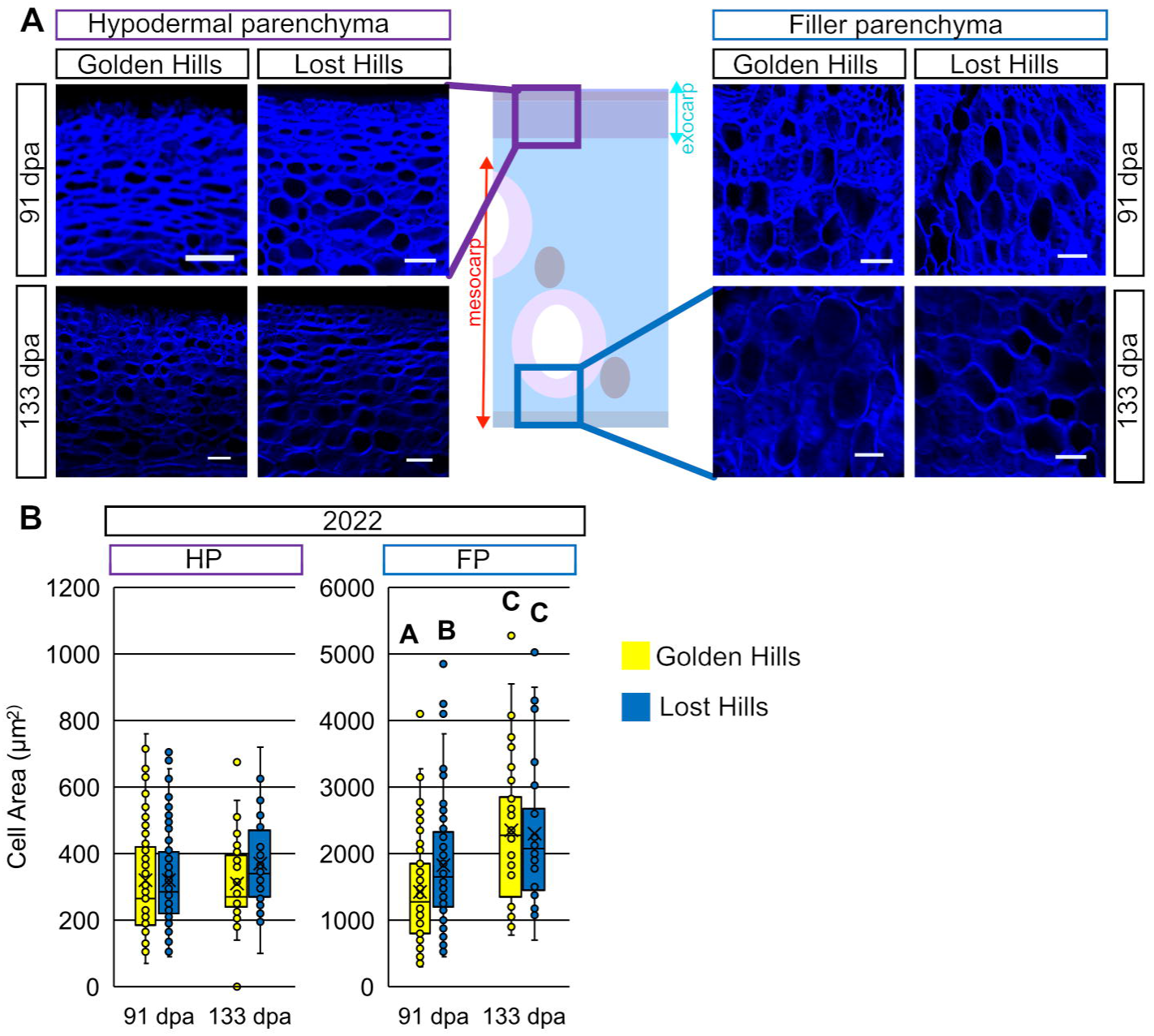
Cell expansion during late-stage hull development. (A-B) Anatomy of hull parenchyma during late-stage hull development. Calcofluor stained hypodermal parenchyma (HP, purple) (A) and filler parenchyma (FP, blue) (B) within the skin tissue of ‘Golden Hills’ and ‘Lost Hills’. Samples were collected at 91 and 133 days post anthesis during the 2022 growing season. Scale bars = 50 µm. Boxes indicate the region of hull sectioned and analyzed. (C) Quantification of HP and FP cross sectional area. N = 30 cells / time point from a minimum of three fruits from three trees. Difference across timepoint is significant for FP, NS for genotype, P < 0.01, two-way ANOVA. Letters indicate difference by Least Square Means with α = 0.05.

The current model of fruit split assumes that an internal expanding force drives the splitting of the fruit skin, led by the turgor pressure from water absorption or changes in the water potential (B.-M. Chang & Keller, 2021; La Spada et al., 2024; Santos et al., 2023). However, questions remain on the nature and contribution of the different interior expansion forces. To test whether pistachio shell split contributes to hull split as an expansion force, we examined the fruits of *P. atlantica,* which only produces fruits with indehiscent shells. The observation of *P. atlantica* fruits with tattered hull suggests that cell expansion, not shell split, plays a more important role in pistachio hull tattering (Sup. Fig. 9).

We reasoned that if FP cell expansion is the main internal expanding force driving hull split, this expansion force would create tension in the hull. In this case, thinner hull tissue would create a mechanically weaker region that is more prone to splitting. To test this hypothesis, we quantified the hull thickness. We identified an interior notch located at the dorsal suture of the hull, resulting in a region that is nearly half as thick as the nonsuture site hull (Fig. 7A-C). Next, we artificially induced freeze-swelling of cells to cause tension in the hull tissue by using liquid nitrogen to snap-freeze intact pistachio fruits. When intact 133 dpa fruit were snap-frozen, the majority of the fruit hull cracked at the dorsal suture, a phenotype similar to the cracked hull observed in the field Fig. 7D). Hulls of younger, intact 105 dpa fruits did not crack as frequently when snap-frozen (Fig. 7D-E). Quantification of the location of hull split for tattered and cracked fruits, showed that indeed the location of hull cracking occurred at the dorsal suture site (Fig. 7B, blue line and 7F), the location with the thinnest hull (Fig. 7A-C), while hull tattering occurred at the nonsuture site (Fig. 7B, white line and 7F). In summary, the data further suggests that a cell expansion force may be the main driver behind hull cracking.

**Fig. 7.**
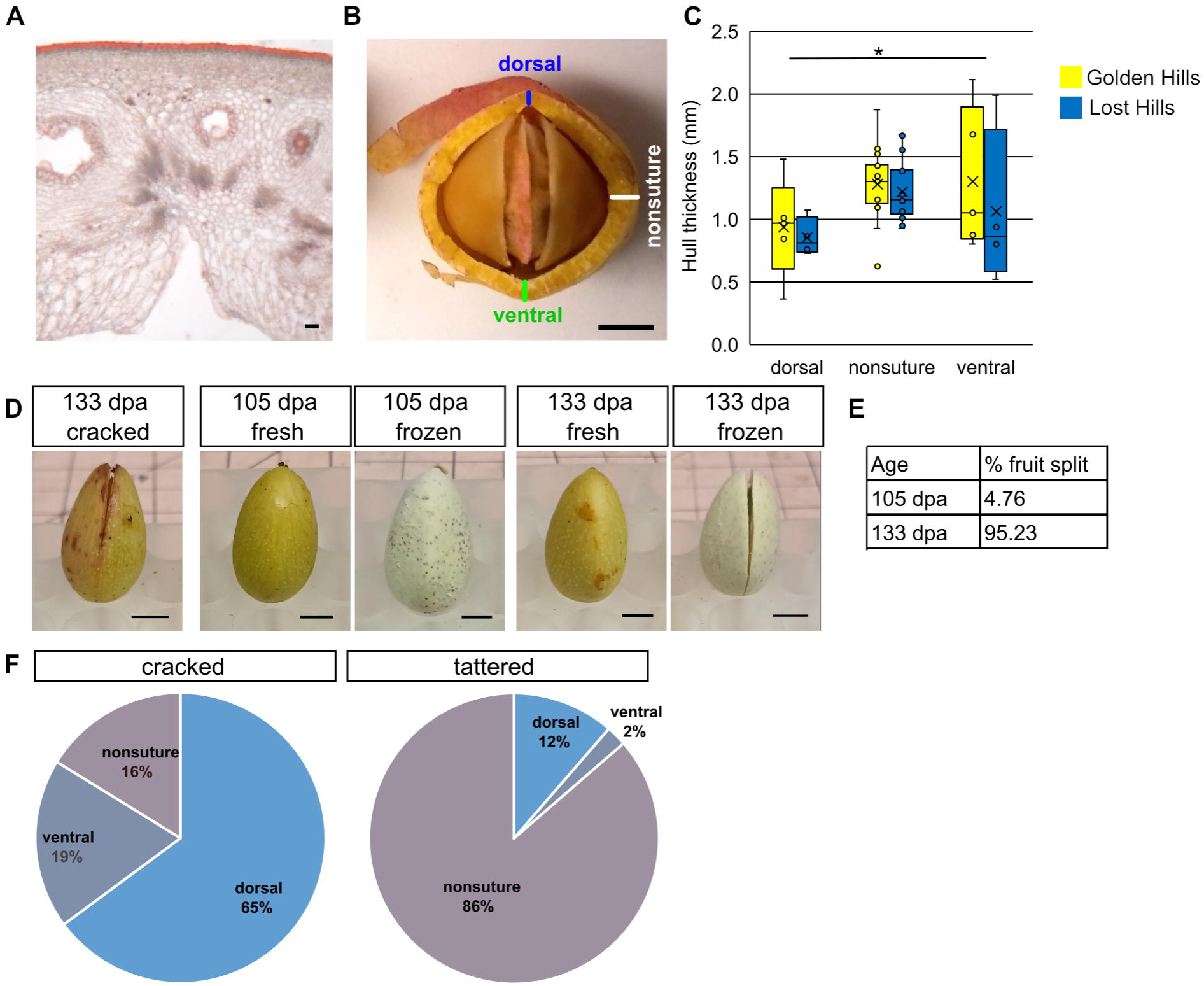
Effect of location-specific hull thickness and simulated cell expansion on hull cracking (A) Anatomy of dorsal suture interior notch at site of cracking initiation. Sudan IV stained hull at dorsal suture shows initiation of tissue split from interior of the fruit. Scale bars = 50 µm. (B) Representative image of 133 days post anthesis (dpa) ’Golden Hills’ fruit cut at median transverse section for measurement of hull thickness at the dorsal suture (blue), nonsuture (white), and ventral suture site (green) showing locations in the hull where split may occur. The lines indicating the thickness measured at each site. Scale bar = 5 mm. (C) Quantification of hull thickness at each site indicated in (B). N = 5-18 fruits from 5-6 trees. P < 0.05 for site, NS for genotype and interaction, two-way ANOVA, NS for Least Square Means post-hoc analysis. (D) Representative image of hull cracking at the dorsal site in field samples and in 133 dpa ‘Golden Hills’ fruit with freezing-induced cell swelling. Similar freezing does not induce cracking at 105 dpa. Scale bars = 5 mm. (E) Percentage of ‘Golden Hills’ fruits that show hull cracking after snap-freezing. N = 21 fruits from 7-8 trees at each timepoint. (F) Location of hull split in cracked and tattered fruits from field samples in 2022. N = 37-44 fruits from 6 trees. P < 0.01, Fisher’s Exact Test.

Finally, cuticle properties such as thickness and integrity have been previously associated with fruit split (B. M. Chang & Keller, 2021a; Dimopoulos et al., 2020; Domínguez et al., 2012; Jiang, Lopez, Jeon, De Freitas, et al., 2019; Knoche et al., 2004; Santos et al., 2023). We examined the cuticles of the different genotypes to test the importance of cuticle in pistachio hull integrity. We did not detect any discernible difference between the cuticles of the cultivars in either thickness or physical appearances over the process of late-stage hull development, between genotypes, and growing seasons 2021 and 2022 (Sup. Fig 10).

### Water status has a significant effect on hull split rate

Environmental factors such as humidity affect fruit split (Correia et al., 2018; La Spada et al., 2024), and likely cell expansion. The difference in ’Golden Hills’ hull phenotypes between 2021 and 2022 at 133 dpa, changing from 95% intact with 5% tattering in 2021 to 79% intact with 16% tattering in 2022 (Fig. 1, Sup. Fig. 2C), suggests that environmental factors may contribute to hull split in pistachio.

To examine whether humidity or irrigation were important drivers behind differences in hull phenotype, we collected fruits from two commercial orchards in 2024, from trees under 50%, 100%, and 150% irrigation treatments. We found that indeed there was a correlation of hull split occurrence with irrigation, where increasing the irrigation from 100 to 150% doubled the frequency of both tattered and cracked hulls (Fig. 8A). In addition, under the same irrigation treatment, prolonged exposure of fruits to humidity significantly increased the hull split percentage, especially of cracked hulls (Fig. 8B). Since fruit cracking is often tested by directly soaking the fruit in water (B.-M. Chang & Keller, 2021; Michailidis et al., 2020), we also tested the pistachio’s response to acute (short term) over-exposure to moisture by spraying fruits from trees in USDA Germplasm Collection at Wolfskill Experimental Orchard (Table 1) with water. Acute exposure to humidity had no obvious effect on the hull split rate (Fig. 8C).

**Fig. 8.**
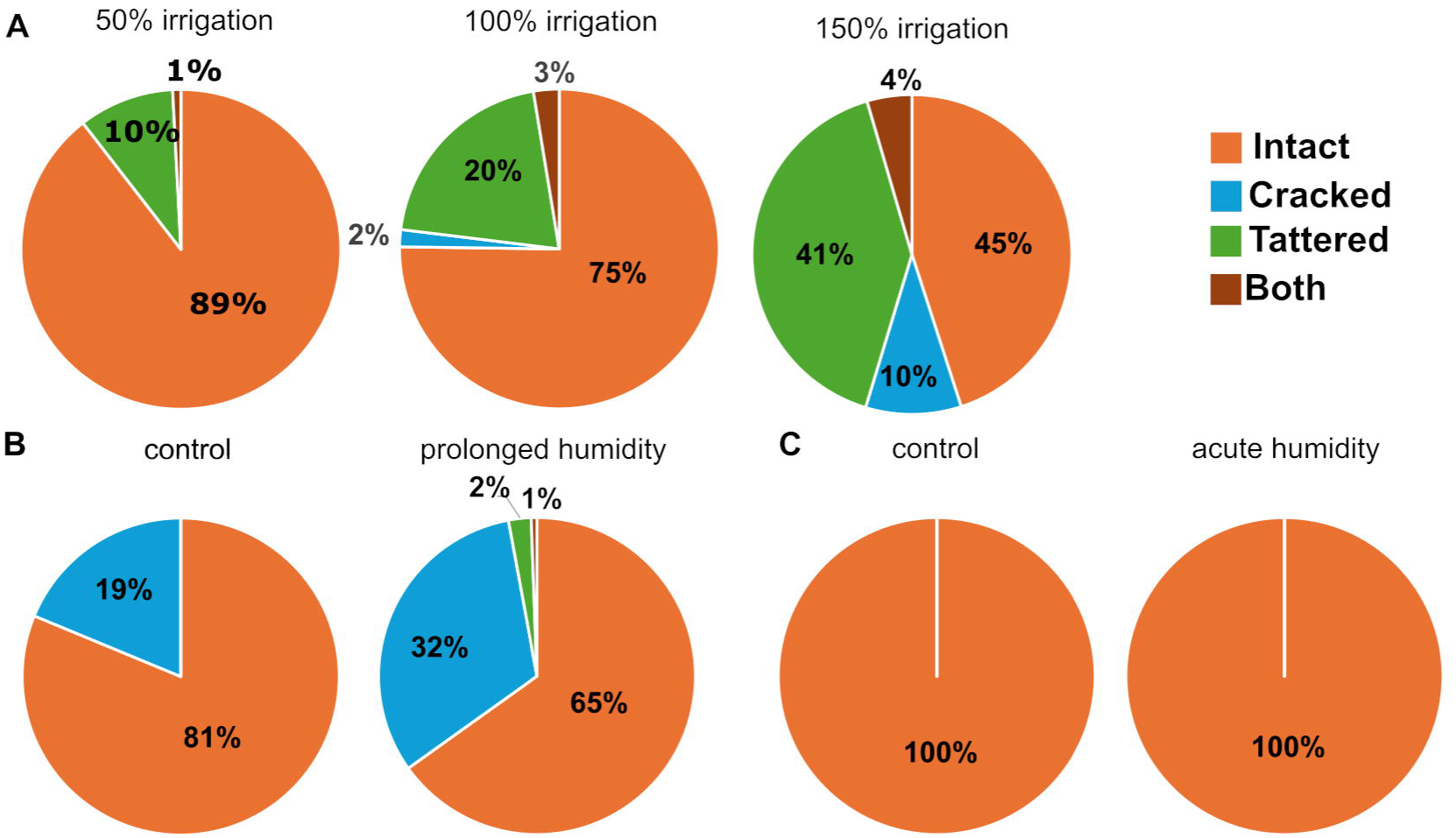
Effect of irrigation and humidity exposure on hull split. (A) Hull phenotype of ‘Golden Hills’ fruits collected from 50%, 100%, and 150% irrigation treatments during the 2024 growing season. N = 331 - 362 fruits from 12 trees per treatment. P < 0.01, Chi-squared test. (B) Hull phenotype of ‘Golden Hills’ fruits under no treatment (control) conditions and after prolonged exposure to moisture. P < 0.01, N = 128 for control, 172 for treated, 8 trees per treatment. Chi-squared test. (C) Hull phenotype of fruits from USDA Germplasm Collection at Wolfskill Experimental Orchard under no treatment (control) conditions and after acute exposure to water. P is NS, N = 104 for control, 159 for treated from 7 trees. Chi-squared test.

To separate out the effect of agricultural management conditions, tree age, rootstock type, and potential genetic variations on hull split, we assayed the hulls of a single tree from the USDA Pistachio Germplasm Collection located at Wolfskill Experimental Orchard from 2022-2024, during which period it was grown under the same orchard management regimen. We observed that the fruits of the D 7 24 tree showed 100% intact hull when harvested between 147-154 dpa in 2022, but showed > 50% hull split in 2023, and 92% intact hulls in 2024 when harvested at the same age (Fig. 9A). This correlated with unseasonal rain during the summer of 2023, which altered the average precipitation, humidity, temperature, and solar radiation during the late fruit developmental stage (Fig. 9B-J). Radiation and relative humidity affect cuticle & cracking (Domínguez et al., 2012; La Spada et al., 2024), and we found that there was indeed a significant increase in dew point and vapor pressure in 2023, with dew point temperature increasing around 2 °C and vapor pressure increasing by 0.3 kPa (Fig. 9B-D). Interestingly, there was no significant difference in air temperature despite the precipitation, and neither soil temperature nor solar radiation correlated with the percentage of hull split (Fig. 9A, E-G). However, there was a change in relative humidity, which was correlated with the total hull split rate across the years (Fig. 9A, H-J).

**Fig. 9.**
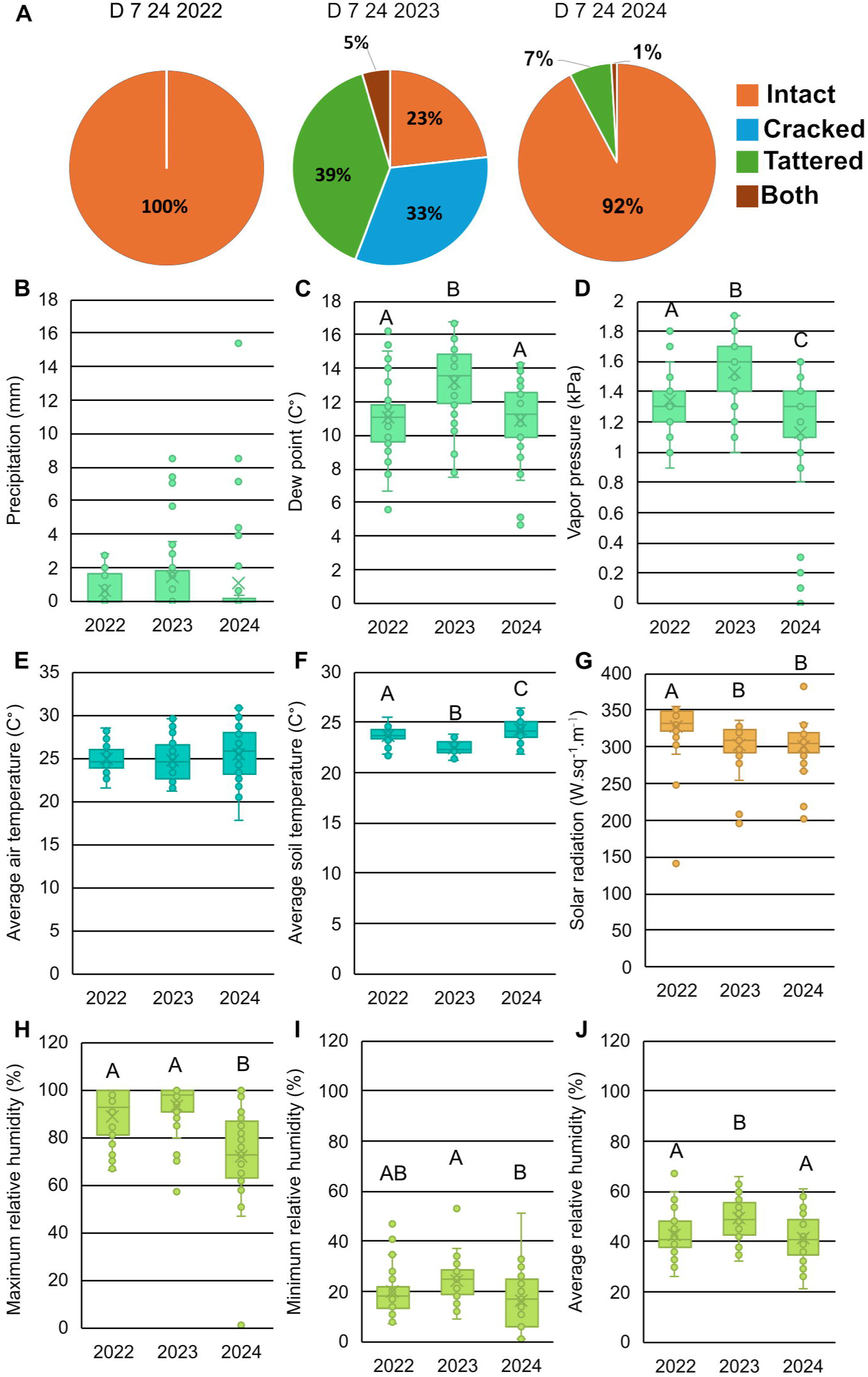
Effect of environmental factors on hull split. (A) Hull phenotype of fruits harvested 147-154 days post anthesis during the 2022-2024 growing seasons from a single *P. vera* tree, D 7 24, in USDA Germplasm Collection at Wolfskill Experimental Orchard. P < 0.01, Chi-squared test. N = 43-219 fruits. (B-J) Average of daily climate data from an onsite weather station at the Wolfskill collection site for 91-133 dpa in 2022-2024. (B) Precipitation, P is NS (C) Dew point temperature, P < 0.01 (D) Vapor pressure, P < 0.01 (E) Average air temperature, P is NS (F) Average soil temperature P < 0.01, (G) Solar radiation, P < 0.01 (H) Maximum relative humidity, P < 0.01 (I) Minimum relative humidity, P < 0.01 (J) Average relative humidity, P < 0.01. N = 41. One-way ANOVA. Letters indicate difference by Least Square Means with α = 0.05.

The ability of the fruit skin to absorb moisture during rain is thought to drive cell expansion and therefore fruit split (B.-M. Chang & Keller, 2021; Cline et al., 1995; La Spada et al., 2024). To test whether there is a potential change in the hull’s ability to absorb moisture during hull ripening due to the formation of microcracks and cell wall modifications, we measured the fruit weight pre- and post-soaking in deionized water from 91 dpa to 142 dpa (Sup. Fig. 11A). Both ’Golden Hills’ and ’Lost Hills’ showed a significant weight increase post-soak at 142 dpa (Sup. Fig. 11A), independent of genotype. Interestingly, we did not detect a significant difference in the ability of the hull to absorb water between the intact hull and the split hull (Sup. Fig. 11B), suggesting that hull integrity does not affect its water uptake properties.

We next tested the effect of the environment on pectin modification and hull integrity by comparing the fluorescence intensity of high and low methylesterified pectin in cultivars we sampled from the 2021-2023 growing seasons (Fig. 10A). We observed that despite the significant variation across years, orchards, trees, and between fruits harvested from the same tree, there was a strong correlation between the fluorescence intensity of low methylesterified pectin in the HP with the frequency of cracked fruits (columns marked in cyan, Fig. 10A, B). There was also a negative correlation between the ratio of HP to FP low methylesterified pectin signal with the percentage of intact fruit (Fig. 10C). The data suggests that the environment and the relative amount of substituted pectin the cell wall, particularly in the HP, are associated with the form of hull split.

**Fig. 10.**
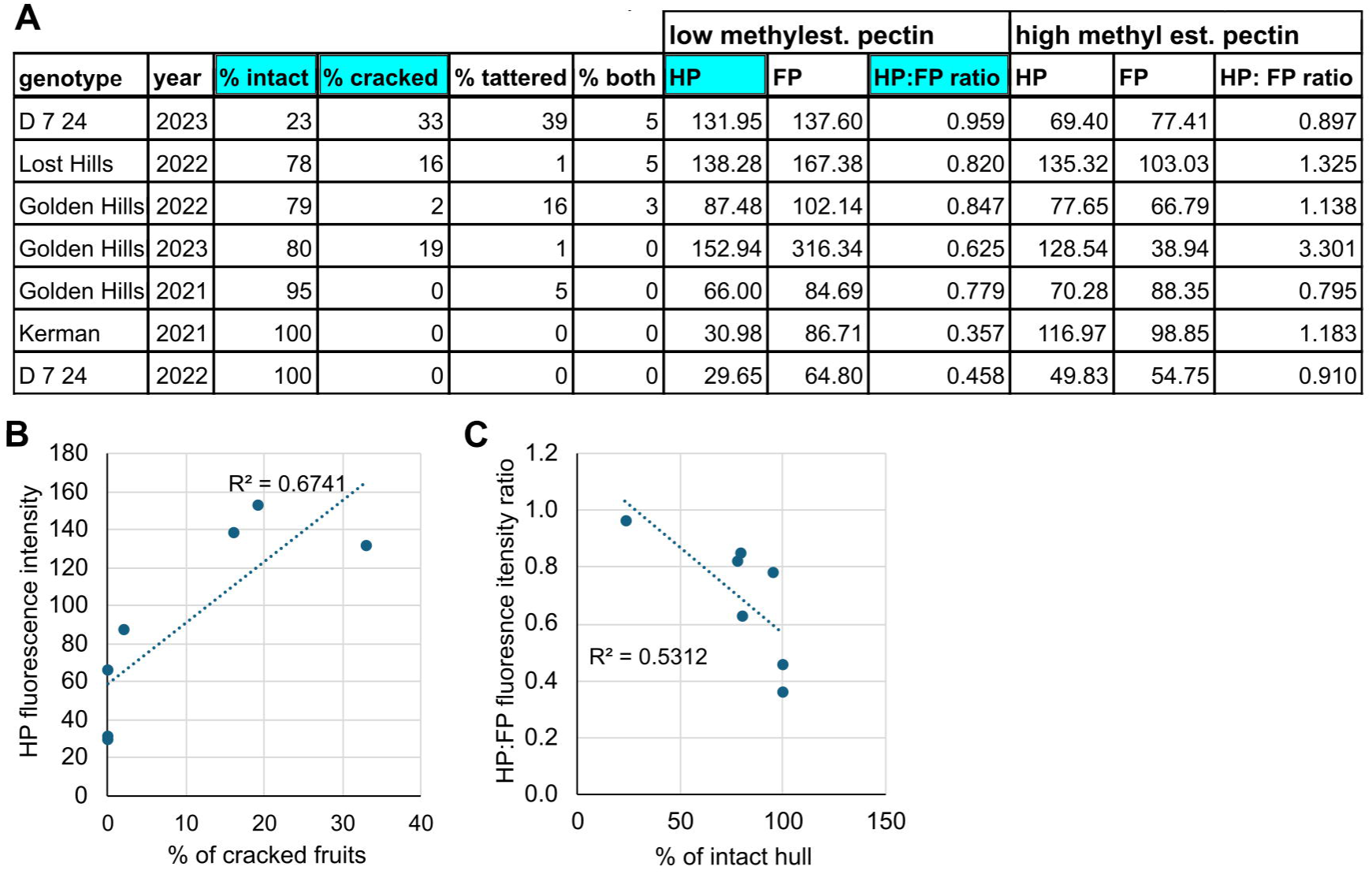
Pectin modification correlates with hull phenotype across growing seasons. (A) Summary of hull integrity, high (JIM7) and low (JIM5) methylesterified pectin signal across different growing seasons in hypodermal parenchyma (HP) and filler parenchyma (FP). Data with strong correlation are in columns indicated in cyan. N = minimum of six fruits sampled from six trees for each immunofluorescence experiment each year. (B) Correlation of JIM5 signal in HP with percentage of cracked hull and (C) correlation of HP:FP ratio of JIM5 fluorescence signal with the percentage of intact hull for data reported in (A).

## Discussion

The anatomy and development of the pistachio hull detailing mesocarp and exocarp tissues, while of economic importance, have not been studied in depth previously. Although recent studies provide a developmental profile of gene expression for the pistachio kernel, shell and hull during ripening (Adaskaveg et al., 2025), the mechanisms behind the exo-mesocarp split in pistachio remain largely unknown. This study provides an anatomical framework of pistachio hull split phenotypes and correlates both transcriptional and *in muro* changes of cell wall polysaccharides across genotypic, developmental, phenotypic, and environmental levels (Fig. 11). Integration of the data shows that the contrast between the expanding FP in the mesocarp and the non-expanding HP in the exocarp may play a critical role in generating the tension necessary for hull split, forming a novel model of fruit split for drupes. In addition, we found that environmental factors may affect cell wall modification and phenotype in a cell-type specific manner (Fig. 11).

**Fig. 11.**
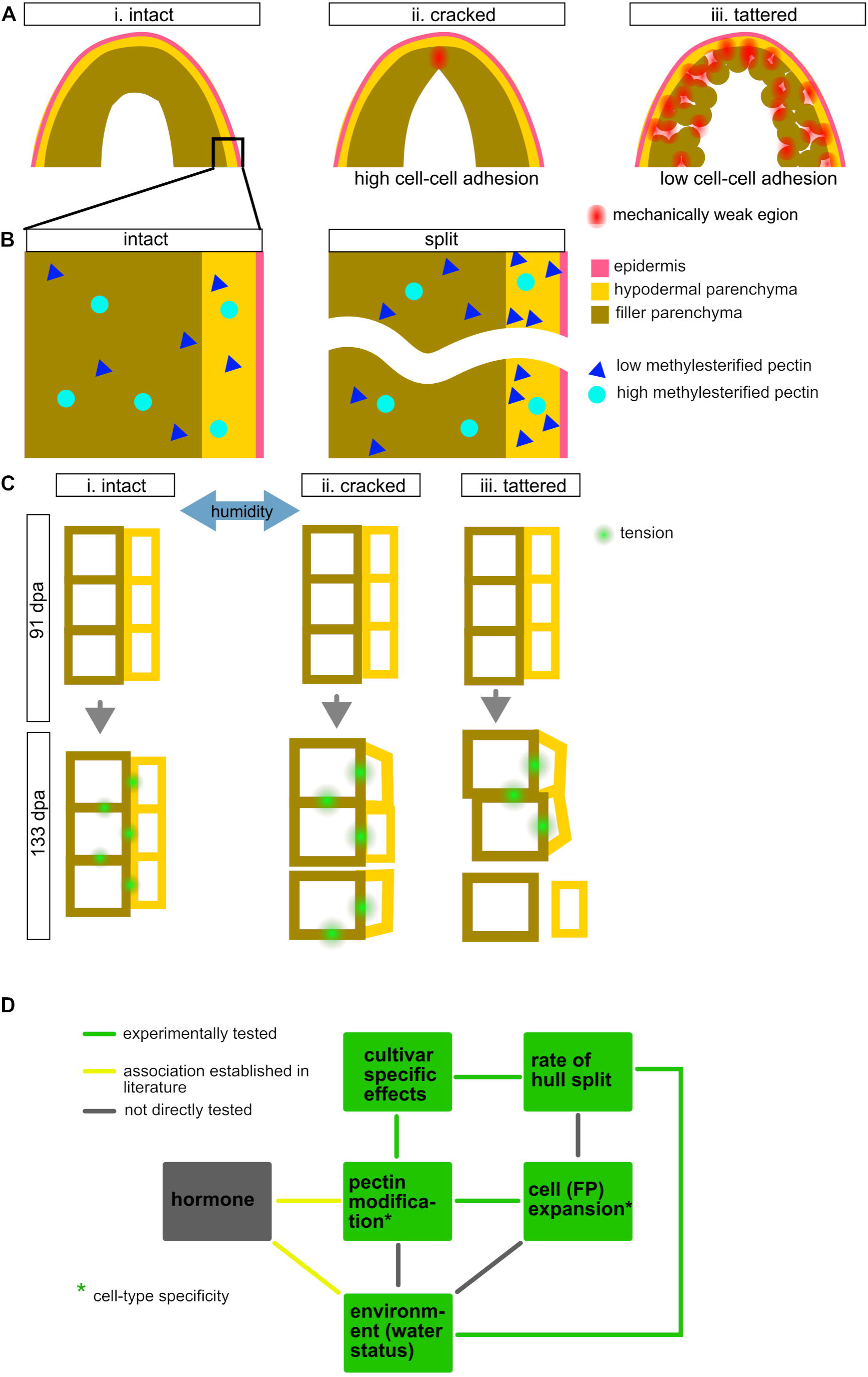
Proposed model of the interacting factors associated with the phenotype and rate of pistachio hull split. (A) Proposed differences in hull mechanical property in intact (i), cracked (ii), and tattered hull (iii) as affected by hull thickness and cell-cell adhesion. (B) Difference in pectin substitution pattern in intact versus split hull. (C) Proposed model of the distribution of tension in intact (i), cracked (ii), and tattered (iii) hull as affected by cell-type specific cell expansion and humidity. (D) Factors (boxes) and interactions (lines) experimentally tested in this study are colored green, established interactions associated with fruit cracking and splitting in other species are colored yellow, factors not associated with hull split tested in this study are colored magenta, while factors and interactions not directly tested are colored grey.

### Pectin remodification occurs during pistachio hull ripening and is associated with cell expansion and hull softening

We have identified PL5 as one of the upregulated genes during late-stage hull development (Fig. 3B). Decrease in cell-cell adhesion, cell wall loosening, and expansion has been previously associated with the pectin lyase (PL) protein family (Huang et al., 2023; Santiago-Doménech et al., 2008; Su et al., 2024; Wang et al., 2019), which supports the role of PL in hull breakdown.

We observed a similar increase in PL expression at our last late-stage hull development time point (Fig. 3B, Sup. Fig. 4), matching what was previously reported (Blanco-Ulate, Barbara et al., 2022), and a correlating decrease in cell-cell adhesion (Fig. 4E). PL5 expression may be higher in intact ‘Golden Hills’ compared to tattered ‘Golden Hills’ at 133 dpa due to the amount of damaged and senescing cells in the latter (Fig. 1B, Sup. Fig. 2B), resulting in tissues that were past the peak of PL activity. Silencing PL has been reported to reduce fruit softening, increase fruit shelf-life, and lead to smaller and more compact skin cells in tomatoes (Ortega-Salazar, 2023; Yang et al., 2017). This suggests that the differences in cell size we observed during pistachio late-stage hull development (Fig. 6) may be attributed to PL activity.

Our immunofluorescence results of low and high methylesterified pectin indicate intense reorganization of the well wall during late-stage pistachio hull development (Fig. 5, Fig. 10). In addition, there is an association between increase in low methylesterified pectin in HP with hull split, particularly of cracked hull (Fig. 10A). Cell wall remodeling, especially of pectin, is a well-established ripening process in many fruits such as sweet cherry, strawberry and tomato, and is associated with fruit softening and fruit split (Basanta et al., 2014; García-Gago et al., 2009; Santiago-Doménech et al., 2008; Santos et al., 2023; Uluisik et al., 2016; Yang et al., 2017). PG and EXPA silenced or double-knockout tomatoes show decrease in low methylesterified pectin and have been associated with firmer fruit (Cantu et al., 2008; Jiang, Lopez, Jeon, de Freitas, et al., 2019) while increase in unesterified pectin is associated with firmness in PG and PL single and double knockout fruits (Ortega-Salazar, 2023; Wang et al., 2019). These studies suggest that the interaction between multiple pectin modifying enzymes and other cell wall metabolic pathways is important for firmness.

Indeed, we observed increased expression of *SNAK1, CHIA, PG, xyloglucan endotransglucosylase/hydrolase, beta-glucosidase* as well as PMEI4 and GAUT3 in our DGE (Fig. 3B, Sup. Table 4-6), indicating multi-level wall remodeling and signaling response that may also overlap with pathogen/ stress response expected during split hull. This leads us to hypothesize that hull softening and splitting may involve mechanisms similar to abscission event in some species such as lupine (*Lupinus luteus*) where increase in low methylesterified pectin can play a role either in cell wall loosening or in plant defense (Wilmowicz et al., 2021). In addition to pectin, cellulose and hemicellulose are known to be involved in fruit softening as well (Shi et al., 2023; Su et al., 2024). Pectin and cellulose modification can act synergistically in changing cell wall elasticity (Altartouri et al., 2019; Su et al., 2024). Changes in pectin can affect cellulose bundling and access of cell modification to cellulose junctions, thus affecting cell wall loosening, elasticity and hydration (Braybrook et al., 2012; Cosgrove, 2016, 2024; Saffer et al., 2023; White et al., 2014). At the same time, PL and PG are known to generate oligogalacturonides, which can function in plant defense signaling (Anderson & Pelloux, 2025; Decreux & Messiaen, 2005; Liu et al., 2024). Hence, the complexity of cell wall modification that is occurring and increased expression of both pectin degradation and biosynthetic gene may explain why we observed an increase in both low and high methylesterified pectin from 91 to 133 dpa (Fig. 5). These cell wall modifications and its interaction with plant defense pathways in split hulls may explain some of the variability we observe in pectin modification (Engelsdorf et al., 2018; Munzert & Engelsdorf, 2025).

A notable finding of this study is that the ratio of low methylesterified pectin in HP versus FP is negatively correlated with hull split (Fig. 10). This can generate a change in cell wall mechanical properties between FP and HP that can lead to either tattering or cracking depending on the location and number of regions of mechanical weakness (Fig. 11A, B). Therefore, the relative levels substituted pectin in different cell layers, in combination with loss of cell-cell adhesion, may create regions with different mechanical properties in the hull, leading to different categories of hull split (Fig. 11A, B).

### Filler parenchyma cell expansion likely contributes to pistachio hull split during late-stage development

Modification in cell walls is necessary for cell expansion (Bou Daher et al., 2018; Braybrook & Jönsson, 2016; Caffall & Mohnen, 2009; Cosgrove, 2000). Snap-freezing only induced cracking at 133 dpa, corresponding to the time period when the hull tissue begins to lose cell-cell adhesion (Fig. 4E). In contrast, 105 dpa hulls have higher cell-cell adhesion, most likely firmer fruit ( Fig 4E) (Adaskaveg et al., 2025), and did not crack upon freezing (Fig. 7D, E). Thus, the changes in cell wall modification as a result of remodeling genes, including the up regulation of the expansin gene family, and the expansion of FP during late-stage fruit development is critical for induced hull cracking (Fig. 3, Sup. Tables 4-6, Fig. 7).

There are two potential sources of internal expansion force in the hull during development: a) the internal force from shell towards the hull as the shell splits and b) the FP cell expansion. The two may act independently or together to increase the tension on the skin. However, the presence of tattered hulls in *P. atlantica,* where the shell is indehiscent led us to conclude that pressure from the shell is not driving hull tattering (Sup. Fig. 9). Thus, we can form a working model of pistachio hull split based on the expansion of FP cells generating the expansion force in the hull (Fig. 6). The lack of similar expansion in HP results in increasing tension in the skin, resembling an over-inflated water balloon. The mechanics of this has been explored in the “theory of shells”, in which fruit split occurs when the skin fails to withstand tension from the interior of the fruit (Bargel et al., 2004; Considine & Brown, 1981). Tracking cell division in HP when compared to FP during late-stage development can be used to explore whether cell wall modification, cell division, or both are contributing to the tension in the skin that leads to cracking.

The distribution of the tension from expanding cells in the FP layers may contribute to the specific phenotype of hull split. We hypothesize that in hulls that have high cell-cell adhesion, the expansion force from the FP is more evenly distributed throughout the hull (Fig. 11C-i). In this scenario, when the hull is experiencing the same amount of tension at all locations, it will tend to break at the region of mechanical weakness, or the thinnest point of the hull, at the dorsal suture (Fig. 7C, 11A-ii). In contrast, if FP cell swelling is occurring in a hull that has low cell-cell adhesion, there will be multiple mechanical weak points throughout the hull (Fig. 11A-iii). These areas and the transition zone where cell senescence is taking place (Fig. 2A-iv) may have lower cell-cell adhesion. In this scenario, if the exocarp is stiff, non-expanding, and mechanically stronger relative to the mesocarp, the expansion force from the FP in the mesocarp may result in the separation of the exocarp from the mesocarp, resulting in the tattering phenotype (Fig. 11A-iii, 11C-iii).

The cuticle can contribute to the tension difference between the exocarp and mesocarp. The properties of the cuticle and its contribution to the stiffness of the fruit skin are involved in grape, tomato, strawberry, and sweet cherry cracking (Dimopoulos et al., 2020; Domínguez et al., 2012; Hurtado & Knoche, 2023; Nadakuduti et al., 2012; Santos et al., 2023). The stiffness of both cuticle and skin is important for grape berry split (B.-M. Chang & Keller, 2021). However, no obvious difference in cuticle between time points and genotypes was observed in our study (Sup. Fig. 10). This suggests that the thicker fruit skin (Fig. 2A) in pistachio and the difference in the relative amount of low methylesterified pectin in the HP and FP layers of split fruits (Fig. 10A-C) may provide the difference in elasticity between skin and interior of the fruit, leading to hull split. Therefore, in our model of fruit split for pistachio (Fig. 11A-C), the HP rather than the cuticle may contribute more to the elastic modulus of the fruit skin.

### Increased irrigation and prolonged humidity promote hull split

Our data indicates that increased irrigation for the entire growing season and prolonged exposure of fruit to humidity can promote hull split (Fig. 8). The effect of humidity on fruit split is consistent with studies in in tomato and other berries, which have shown that rainfall and humidity are involved in fruit splitting and cracking (Cline et al., 1995; Correia et al., 2018; Domínguez et al., 2012; Haokip et al., 2020; La Spada et al., 2024; Santos et al., 2023).

Environmental effects can also help explain why ‘Golden Hills’ showed high rates of hull integrity in 2021 (Sup. Fig. 2), but high rate of tattering in 2022 and 2023 growing seasons under normal irrigation (Fig. 1B, Sup. Fig. 2, Fig. 8B). This is further supported by the phenotype of the tree D 7 24 in Wolfskill Experimental Orchard (Fig. 9A). D 7 24, under the same irrigation regime, showed high integrity in 2022 and 2024, but high cracking and tattering in 2023 (Fig. 9A), and the hull split frequency of D 7 24 correlated strongly with air humidity (Fig. 9C, D, J). However, maximum acute exposure to moisture, similar to assays commonly performed in grapes and sweet cherries (B.-M. Chang & Keller, 2021; Michailidis et al., 2020), did not increase split rate (Fig. 8C), while prolonged exposure did (Fig. 8B), suggesting that the length of time that the tension in the hull is maintained is more important than the intensity. Future work is needed to more directly assess the hull skin tension and osmotic potential of the hull tissue under different humidity and irrigation conditions, as well as to determine if there is strong correlation between cell turgor pressure, cell wall swelling, and hull split under different humidity and irrigation conditions.

In conclusion, in our study we have identified two main parenchyma cell types key to hull split. Cell wall modification, especially of pectin, occurs during late-stage pistachio hull development, and this modification is associated with hull softening and loss of cell-cell adhesion. We established the complex interactions between phenotype and environment that occur at the molecular and cellular level and identified candidate genes for cell wall modification and cell expansion. Furthermore, we showed that increased irrigation and prolonged exposure to humidity promotes hull split, suggesting a possible link between water status and changes at the cellular level. Together, we have built a model where cell-type specific expansion forms the main driving force behind hull split, while the changes in cell wall modification and cell adhesion may determine the amount of cell expansion and category of hull split.

## Supplemental data

Table S1. USDA Germplasm ID from Wolfskill Experimental Orchard of trees used in this study in various assays.

Table S2. Primers used in qRT-PCR.

Fig. S1. Heatmap of RNAseq read count of actin, *EF1a*, and *GAPDH* gene families commonly used as reference genes in qRT-PCR of plant tissues.

Fig. S2. Additional characterization of hull split phenotypes.

Table S3. Statistical analysis of the RNAseq read quality.

Fig. S3. Differential gene expression analysis summary of RNAseq.

Fig. S4. qRT-PCR of genes of interest identified in RNAseq.

Table S4. Top ten genotype-specific candidate genes identified in DGE analyses comparing ‘Golden Hills’ and ‘Lost Hills’ at 91 and 133 dpa.

Table S5. Top ten phenotype-specific candidate genes identified in DGE analyses comparing intact, cracked, and tattered hull at 133 dpa.

Table S6. Top ten candidate genes for late-stage hull development identified in DGE analyses comparing 91 dpa hull against intact, cracked, and tattered hull at 133 dpa.

Fig. S5. RNAseq Gene Ontology and KEGG enrichment analyses.

Fig. S6. Immunofluorescence analyses for low methylesterified pectin (JIM5) and high methylesterified pectin (JIM7).

Fig. S7. Negative control for antibodies and stains used.

Fig. S8. Estimated cell size of hull hypodermal and filler parenchyma for fruits collected in 2021.

Fig. S9. Hull split in *Pistacia atlantica*.

Fig. S10. Cuticle thickness of pistachio cultivars during late-stage fruit development.

Fig. S11. Hull tissue absorption during late-stage fruit development.

## Acknowledgement

The authors would like to thank Professor Judy Jernstedt for advice on anatomical identification and sectioning, and acknowledge the use of Zeiss 980 confocal microscope made available through the National Institute of Health grant S10D026702 and the technical Director of the Light Microscopy Core Dr. Wilkop for guidance on image acquisition and processing.

## Contributions

S.Z., G. D., and P. G., conceived the research and designed the experiments. S. Z., S. E., A. C., M. W., A. B., P. G., K. M. B., P. T., and R. L. performed the wet lab experiments. C. L., J.A.A., Y.W., and G. M. performed the assembly and annotation of the reference genome. S. Z. and G. D. wrote the manuscript. S. Z., J.A.A. and BB-U, and G. D. contributed to the review and editing. All authors discussed the results. All authors have read and agreed to the published version of the manuscript.

## Data availability

The RNAseq data underlying this article are available in the Gene Expression Omnibus Database at https://www.ncbi.nlm.nih.gov/geo/ and can be accessed with the accession number GSE300184.

